# Sox9 and Nfi transcription factors regulate the timing of neurogenesis and ependymal maturation in dopamine progenitors

**DOI:** 10.1101/2024.11.12.623209

**Authors:** Laura Lahti, Nikolaos Volakakis, Linda Gillberg, Behzad Yaghmaeian Salmani, Katarína Tiklová, Nigel Kee, Hilda Lundén-Miguel, Michael Piper, Richard Gronostajski, Thomas Perlmann

## Abstract

Correct timing of neurogenesis is critical for both generating the correct number and subtypes of glia and neurons in the embryo, as well as preventing tumours and the depletion of stem cell pools in the adults. Here we analyse how the midbrain dopamine neuron (mDA) progenitors transition into cell cycle arrest (G0) and begin to mature into ependymal cells. The comparison of mDA progenitors from different embryonic stages revealed the upregulation of *Nfi* and *Sox9* transcription factors during development. Their conditional inactivation in the early embryonic midbrain leads to delayed G0 entry and ependymal maturation, reduced gliogenesis, and increased generation of neurons, including mDA neurons. In contrast, their inactivation in late embryogenesis does not result in mitotic re-entry, suggesting that these factors are necessary for the G0 induction, but not for its maintenance. Our characterisation of mDA-progenitor-derived adult ependymal cells by single-cell RNA sequencing and histology show that they both retain several progenitor features but also secrete neuropeptides and contact neighbouring cells and blood vessels, indicating that these cells may form a novel part of the circumventricular organ system.

## Introduction

Dopamine, one of the brain’s neurotransmitters, is pivotal in orchestrating motor behaviors, cognition, memory, and reward mechanisms. Dopamine transmission is facilitated by a diverse population of midbrain dopamine (mDA) neurons that exhibit distinct innervation patterns and functions (Yaghmaeian et al., 2024; Poulin et al., 2020; Garritsen et al., 2023; Azcorra et al., 2023).

How mDA neurons are generated during embryonic development has attracted significant attention, mainly due to their importance in Parkinson’s disease (PD). This neurodegenerative disease is characterised by the degeneration of mDA neurons, resulting in motor complications. Molecular understanding of mDA neuron development has enabled the generation of stem cell-derived mDA neuron precursors *in vitro*. Several on-going clinical trials test transplants of these *in vitro* -generated cells, aiming at restoring DA signaling in PD patients (Kirkeby et al., 2023; Piao et al., 2021; Doi et al., 2020; Barker and Björklund, 2023).

Another – currently hypothetical – approach to replace mDA neurons lost in PD could be the stimulation of endogenous brain repair. For instance, the loss of dopamine-mediated feedback inhibition to neighbouring astroependymal cells in salamanders can induce re-entry into the cell cycle leading to generation of new mDA neurons (Berg et al., 2011). Such ability appears absent in the mammalian midbrain, which does not harbour neural stem cells in the adult. Instead, neural stem cells reside in few restricted areas in the adult mammalian brain: in the subventricular zone (SVZ) in the cortex, and in the hippocampus, where they reside in a state of non-permanent cell cycle arrest (quiescence) (Urbán et al., 2019).

Adult neural stem cells are formed during embryogenesis, when some neural progenitors are specified to enter quiescence and form the stem cell reserve (Fuentealba et al., 2015; Furutachi et al., 2015; Morales and Mira, 2019). After generating neurons and glia, the remaining progenitors exit the cell cycle and differentiate into multi-ciliated ependymal cells. These cells line the brain ventricles and the spinal cord central canal and contribute to the maintenance of CSF homeostasis as well as to neuroblast migration (Ji et al., 2022). In the spinal cord, ependymal cells can proliferate *in vivo* and contribute to glial scar formation after injury (Meletis et al., 2008; Barnabe-Heider et al., 2010, Sabelström et al., 2013; Ren et al., 2017), but in the brain, the regenerative capacity of the ependymal cells remains unclear (Chiasson et al., 1999; Spassky et al., 2005; Zhang et al., 2007; Carlén et al., 2009; Shah et al., 2018). Also progenitors in the midbrain floor plate, which generate mDA neurons between embryonic day (E)10 and E14 (Bayer et al., 1995), become ependymal cells later. However, these cells appear to retain the expression of mDA progenitor marker *Lmx1a* even in adult (Hedlund et al., 2016). This raises the question of whether the ependymal cells in different brain regions have dissimilar properties and functions.

Understanding the induction of mitotic exit in neural progenitors and the maintenance of G0 in the derivatives of neural progenitors is important for several reasons. First, dysregulation of the cell cycle exit of radial glial cells leads to an imbalance in the types and amounts of cells generated, which can give rise to various neurodevelopmental disorders such as autism (Kaushik and Zarbalis, 2016), schizophrenia (Mao et al., 2009), and ADHD (Dark et al., 2018). In addition, incomplete maturation of postmitotic radial glia into ependymal cells can lead to hydrocephalus (Jiménez et al., 2001; Park et al., 2016, Hou et al., 2023). Second, the failure to maintain quiescence in the adult neural stem cells will lead to the depletion of the stem cell pool (Velthoven and Rando, 2019) and can contribute to neurodegenerative disorders (Joseph et al., 2020). Indeed, a substantial effort has been made to identify signaling pathways and transcription factors responsible the maintenance of stem cell quiescence (Blasco-Camarro and Fariñas, 2023). Although many of them are shared between the embryonic and adult neural stem cells, some serve somewhat different or even opposing functions (Urbán and Guillemot, 2014). For example, transcription factors belonging to Nuclear Factor I -family (Nfi) promote neurogenesis in the embryonic telencephalon, but inhibit it to maintain neural stem cells in the adult (Harris et al., 2015).

In this work, we have analysed the regulation of the timing of neurogenesis and mitotic exit in the midbrain, focusing on the mDA-neuron-generating floor plate. We characterised transcriptional differences of floor plate radial glia from different embryonic stages – from the active phases of mDA neurogenesis to the ependymal maturation. Together, our data revealed a molecular basis for the tight temporal control of mitotic exit as well as neuro- and gliogenesis in the midbrain, including the region generating mDA neurons.

## Results

### *Nfi* transcription factors and *Sox9* are upregulated in late stage mDA progenitors

In order to understand the timing of mitotic exit in the mDA progenitors, we first asked whether they produce glia after having generated mDA neurons. For this, we administrated tamoxifen at E9.5 to embryos carrying *Lmx1aCreERT2* and *R26*^*TrapCherry*^ (TrapC) reporter. The tamoxifen treatment activated Cre in mDA progenitors, labelling them and all their progeny with red fluorescent protein (RFP) mCherry (**Supp. Fig. 1A**). We could verify that RFP+ cells in the adult midbrain had neuronal morphology and expressed NeuN. None were GFAP^+^ nor Olig2^+^ (**Supp. Fig. 1B,C**). This indicated that after generating mDA neurons, the mDA progenitors lack a gliogenic phase and likely exit mitosis.

To understand the transcriptional changes during the transition into cell cycle exit, we re-analysed our previously generated single-cell RNA sequencing (scRNA-seq) dataset of mDA progenitors (Kee et al., 2017). This revealed clusters containing either “early” (E10-11) and “late” (E12-13) *Lmx1a*^*+*^ *Corin*^*+*^ cells that expressed progenitor markers *Hes1, Nestin*, and *Mki67*, indicating that these cells were mDA progenitors (**Supp. Fig. 2A-C**). The “late” cluster cells expressed reduced levels of *Ccnd1*, suggesting some of these cells may have already exited the cell cycle. The single-cell differential expression (SCDE) analysis between the “early” and “late” clusters yielded lists of genes either down- or upregulated in “late” mDA progenitors (**Supp. Fig. 2D,E, Supp. Table 1**). The downregulated genes were involved in cell cycle regulation, while the list of upregulated genes contained two transcription factors previously associated with the regulation of stem cell and progenitor quiescence, *Nfib* and *Sox9* (Chang et al., 2013; Scott et al., 2010, Kadaja et al., 2014, Clark et al., 2019).

To complement the scRNAseq dataset from later embryonic stages, Lmx1a^CreERT2^-labelled floor plate cells were collected from E11, E13, and E18 embryos for RNA sequencing by laser capture microdissection, LCM (**Fig. 1A, Supp. Table 1**). Comparing the differentially expressed genes between E13 and E11 revealed upregulation of *Nfib* and *Nfix* and, to a lesser extent, *Nfia* (**Fig. 1B**). By checking all known *Nfi* factors and *Sox9* in the scRNAseq dataset, *Sox9* was already present in the “early” mDA progenitor cluster, as was *Nfia* to some extent (**Supp. Fig. 2F**). In contrast, *Nfib* and *Nfix* were more enriched in the “late” cluster, whereas *Nfic* was present only in few cells and was neither significantly upregulated in our E13 vs E11 LCM-RNAseq analysis.

**Fig. 1.**
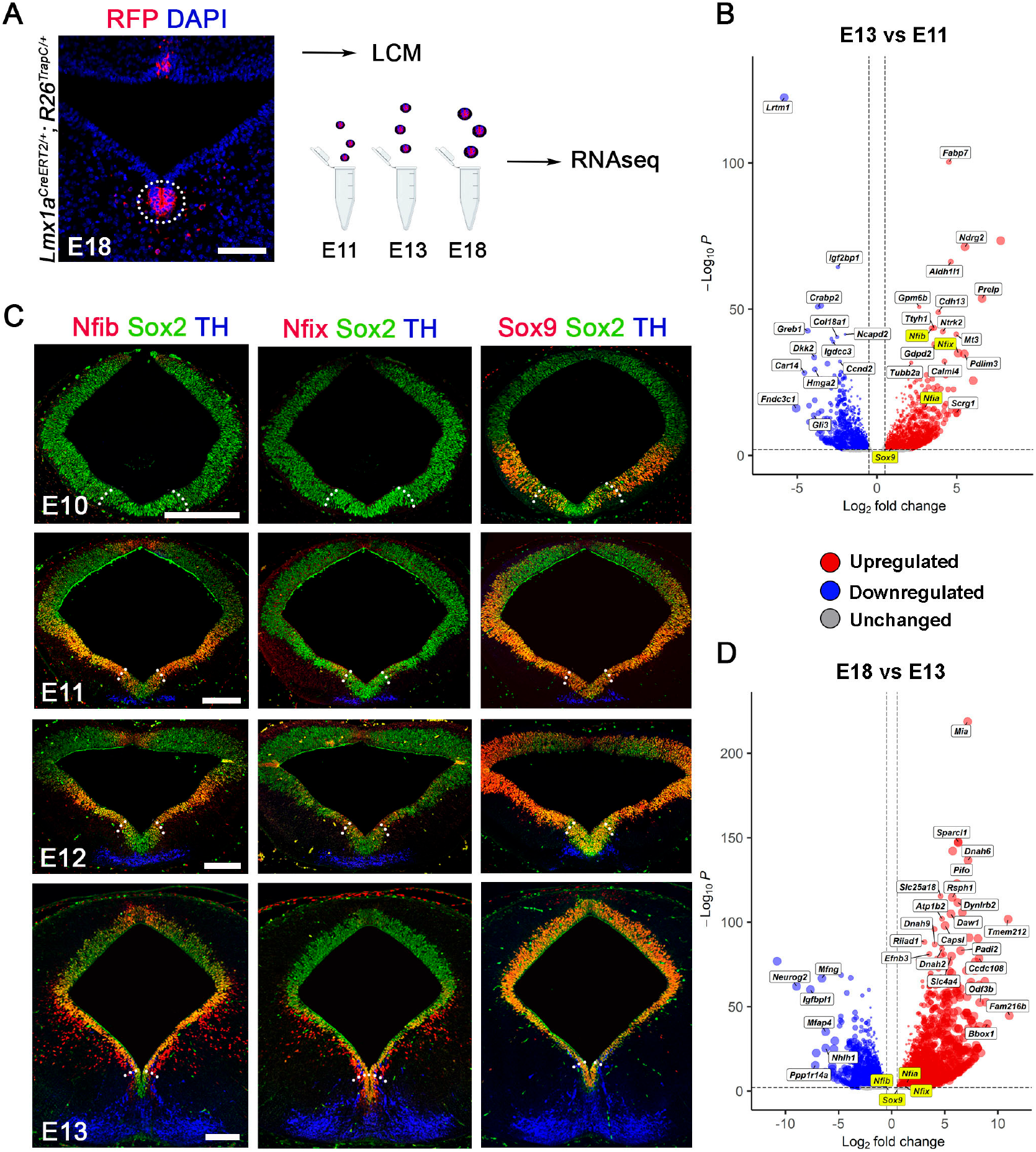
Nfi and Sox9 transcription factors are upregulated in the mDA progenitors towards the end of mDA neurogenesis. **A)** Schematic overview of lasercapture microdissection (LCM) and bulk RNA sequencing of mDA progenitors. The dotted circle indicates the area collected. **B)** Volcano plot visualising selected variable genes (4939) enriched in mDA progenitors at E13 (red) vs E11 (blue). **C)** Immunohistochemical detection of of Nfib, Nfix, and Sox9 in the midbrain. The mDA progenitor domain is indicated by dotted lines. **D)** Volcano plot visualizing selected variable genes enriched in the E18 (red) vs E13 (blue) DA progenitor domain (8075). Scalebars 100 μm in A and 200 μm in C.

Immunohistochemical analysis of these factors in the embryonic midbrain showed that their expression dynamics followed an approximately similar pattern with signal first detected in the ventrolateral ventricular zone (VZ), followed by upregulation in the mDA progenitors of the floor plate. At E10, when the first mDA progenitors become post-mitotic, Sox9 started to be expressed in the ventral midbrain, where *Nfis* were still absent (**Fig. 1C, Supp. Fig. 3B**). Upregulation of *Nfia* and *Nfib* began ventrolaterally at E11 and Nfix at E12. By E13, when mDA neurogenesis begins to cease, the floor plate expressed *Sox9, Nfix*, and *Nfia*, whereas *Nfib* expression became apparent by E14 (**Supp. Fig. 3A**). In contrast, *Nfic* was restricted to roof plate (**Supp. Fig. 3C**). Supporting these results, the E18 versus E13 comparison of the dataset showed no expression changes of these factors, suggesting a plateau in G0-state mDA progenitors (**Fig. 1D**). These results show that *Nfib, Nfix, Nfia*, and *Sox9* become gradually upregulated in the midbrain VZ during the neurogenic period and that by the end of mDA neurogenesis, they are all expressed in the floor plate.

### *Nfib, Nfix*, and *Sox9* are required for the correct timing of mitotic exit in midbrain VZ

These observations led us to speculate that Sox9 and Nfi transcription factors might regulate the transition from proliferation to G0 in the midbrain neuronal progenitors, and that the inactivation of these factors might delay or inhibit this transition.

To analyse this, we conditionally inactivated these alleles in the midbrain by recombining the floxed alleles with En1^Cre^, which is expressed in the midbrain and anterior hindbrain from E8.5 onwards (Kimmel et al., 2000; Sgaier et al., 2005). As *Nfia* was already strongly expressed in all mDA progenitors already at E11, it may be less critical for regulating their mitotic exit and was thus excluded from these functional studies. Although *Sox9* displayed similar expression pattern, being even earlier expressed than *Nfia*, we chose to include this factor for two reasons: first, it had been shown to act upstream of *Nfia* in the developing spinal cord (Kang et al., 2012), suggesting that it might regulate Nfi transcription factors also in the midbrain, and second, its conditional inactivation in the embryonic cerebellum results in prolonged neurogenesis (Vong et al., 2015).

We generated both single mutants and different combinations of these alleles, noting that they did not affect the expression of each other or *Nfia* (**Supp. Fig. 3E**). These results also showed that in the midbrain, *Sox9* was not required for the expression of *Nfis*. We detected no apparent midbrain phenotype in *Nfib*^*ck*o^ or *Nfix*^*cko*^ single mutant embryos (data not shown), and they are not described further in this study. Instead, we focused on *Sox9*^*cko*^, *NfixNfib*^*dcko*^ and *Sox9NfibNfix*^*tck*o^ mutants.

To analyse the proliferative state of the midbrain VZ in these mutants, we pulsed them with BrdU for 30 minutes before tissue collection at different embryonic stages and compared the proportion of BrdU^+^Sox2^+^ cells to all Sox2^+^ cells in the VZ. As these factors had shown different expression dynamics in the dorso-ventral domains of the midbrain (**Fig. 1C**), we analysed separately the most dorsal tip (roof plate), the most ventral tip (mDA-generating Lmx1a^+^ floor plate), and the dorsolateral and ventrolateral midbrain.

As *Nfib, Nfix* and *Sox9* were gradually upregulated in the midbrain (**Fig. 1C**), the lack of major differences in BrdU uptake between the genotypes at earliest stages analysed was unsurprising (E11 and E12; **Supp. Fig. 4A**). In contrast, a few days later numerous BrdU^+^ nuclei could be detected in the VZ of all mutants (**Fig. 2A**). Quantifying the signal across the stages showed the most drastic increase in BrdU uptake in *Sox9NfibNfix*^*tcko*^ VZ (**Fig. 2B**). The requirement of these factors differed across the dorsoventral domains, with *Sox9*^*cko*^ showing a significantly higher proportion of cycling cells in the ventrolateral domain compared to *NfibNfix*^*dck*o^ (**Fig. 2B**). These results indicate that the loss of *Sox9, Nfib*, and *Nfix* keeps the neural progenitors longer in the cell cycle and postpones the entry into G0.

**Fig. 2.**
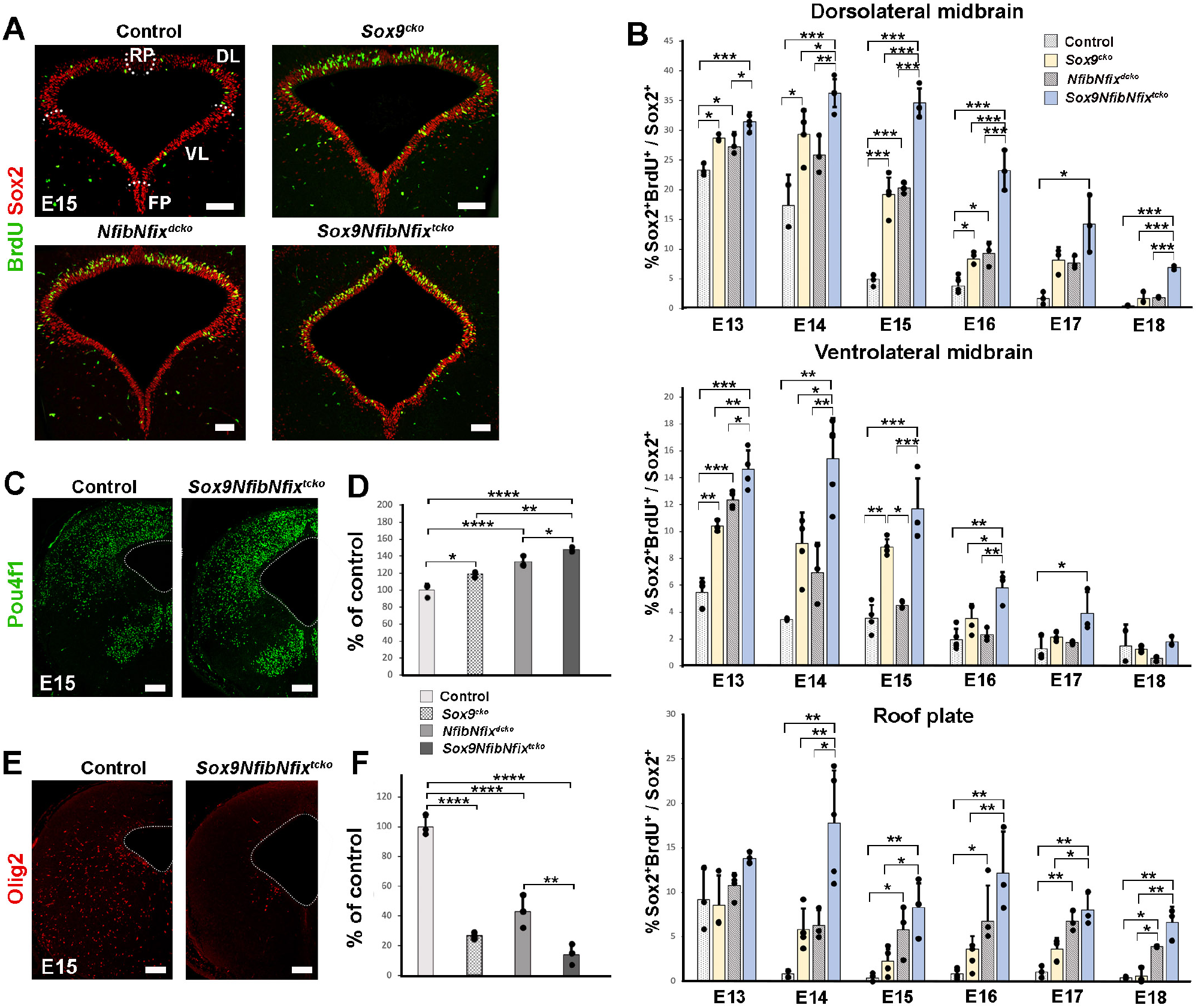
Loss of *Sox9* and *Nfi* transcription factors delays mitotic exit in neural progenitors. **A)** IHC staining of E15 midbrains after a 30-minute BrdU-pulse. **B)** Quantification of BrdU uptake. The domains are indicated by dotted lines in A. **C)** IHC staining for Pou4f1^+^ glutamatergic neurons with quantification in **D** (n=3/genotype). **E)** IHC for an Olig2^+^ oligodendrocyte precursors with quantification in **F** (n=3/genotype). The ventricle is visualized with the dotted line in C and E. FP, floor plate, VL, ventrolateral midbrain, DL, dorsolateral midbrain, RP, roof plate. Data shown as mean (SD). Only comparisons with p-value <0.05 are indicated. * p<0.05, ** p<0.01, *** p<0.001. One-way ANOVA with Tukey post hoc. The information of technical and biological replicates is found in Supp. table 5. Scalebars 100 μm in A and 200 μm in C and E.

The prolonged proliferative state might also disturb or postpone the maturation of progenitors into ependymal cells. We analysed this by immunohistochemical staining for FoxJ1, a regulator of ciliogenesis in ependymal cells (Jacquet et al., 2009), and S100b, a marker for mature ependymal cells (Vives et al., 2003). We could detect fewer FoxJ1^+^ and S100b^+^ cells in all mutants at E18, with an almost complete absence of S100b^+^ cells in *Sox9NfibNfix*^*tcko*^ VZ (**Supp. Fig. 4B,C**). This suggests that - consistent with the results from the *Nfix*^*-/-*^ cortex (Harkins et al., 2022) - the loss of *Nfib, Nfix* and *Sox9* delays or even prevents full ependymal maturation in the midbrain.

### The loss of *Nfib, Nfix*, and *Sox9* leads to prolonged neurogenesis and impaired gliogenesis in the midbrain

As cell cycle exit is linked to neurogenesis and gliogenesis, we investigated whether the increased proportion of cycling cells in *Nfib/x-Sox9* mutant VZ had any effect on these processes. The ventrolateral and dorsolateral progenitor domains of the midbrain generate both glutamatergic, GABAergic, and some cholinergic, neurons (Lahti et al., 2013). At E15 the number of Pou4f1^+^ glutamatergic neurons was increased in the midbrain in all mutants compared to the controls (**Fig. 2C,D**). An opposite effect was seen in the production of Olig2^+^ oligodendrocyte precursors, with the highest reduction in *Sox9NfibNfix*^*tcko*^ embryos (**Fig. 2E,F**). Furthermore, all mutants seemed to have fewer Aldh1l1^+^ cell somata in the midbrain marginal zone, indicating that the generation of astrocytes was also affected (**Supp. Fig. 4D**).

These results indicate that similar to what has been shown in other tissues (Stolt et al., 2003; Deneen et al., 2006; Kang et al., 2012, Clark et al., 2019), Nfib, Nfix and Sox9 both promote gliogenesis in the midbrain, and that by inducing mitotic exit in the progenitors, they also control the extent of neurogenesis.

### Prolonged and increased generation of dopamine neurons in the absence of *Nfib, Nfix*, and *Sox9*

We then turned our attention to the mDA neuron -generating floor plate. Similar to more dorsal midbrain, floor plate progenitors showed increased BrdU uptake in all mutants compared to the controls, with the strongest effect in the *Sox9NfibNfix*^*tcko*^ mutants (**Fig. 3A,B**). By E14, when mDA neurogenesis is normally completed, only few individual BrdU^+^ Sox2^+^ cells were detected in the floor plate of the control embryos. In contrast, the mutants showed higher BrdU uptake until E16 (**Fig. 3B**).

**Fig. 3.**
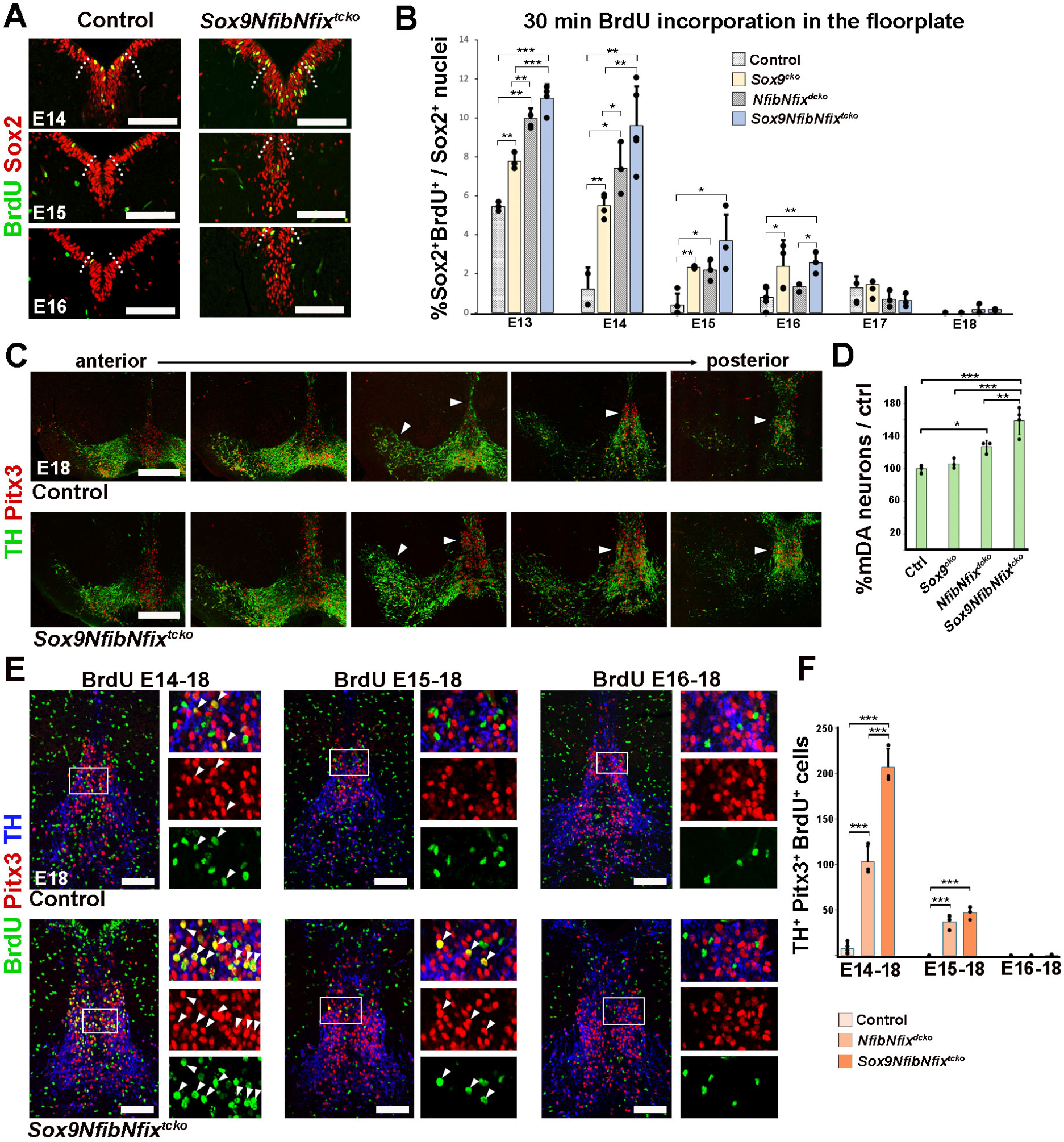
Increased and prolonged mDA neurogenesis in *Nfib/x-Sox9* compound mutants. **A)** IHC staining of ventral midbrains after a 30-minute BrdU-pulse. The boundaries of *Lmx1a*^*+*^ floor plate indicated with dotted lines. **B)** Quantification of BrdU+ cells in the floor plate. **C)** IHC stainings of mDA neurons from anterior to posterior. Arrowheads indicate regions with highest increase in mDA neurons in mutants. **D)** Quantification of mDA neurons at E18 (Ctrl and *Sox9NfibNfix*^*tcko*^ n=4, *Sox9*^*cko*^ and *NfibNfix*^*dck*o^ n=3). **E)** IHC of mDA neurons from different BrdU-labelling schemes, images from caudal VTA. Squares indicate the regions for high magnification images. Arrowheads point to BrdU^+^Pitx3^+^TH^+^ cells. Quantification in **F** (ctrl n=5, others n=3). Data shown as mean (SD). Only comparisons with p-value <0.05 are indicated. * p<0.05, ** p<0.01, *** p<0.001. One-way ANOVA with Tukey post hoc. The information of technical and biological replicates is found in Supp. table 5. Scalebars 100 μm in A,E and 200 μm in C.

Furthermore, neurogenic genes *Neurog2, Ascl1*, and *Dll1* were upregulated in the E14 *Sox9NfibNfix*^*tcko*^ ventral midbrain compared to the control, indicating prolonged neurogenesis (**Supp. Fig. 5A**). Hes5, a marker for both neural and glial progenitors, showed only upregulation in more lateral midbrain of the mutants and was absent in the non-gliogenic floor plate. Cell cycle G1-S transition regulator CyclinD1, and PCNA, another marker for cycling cells, appeared also upregulated in E15 *Sox9NfibNfix*^*tcko*^ ventral midbrain (**Supp. Fig. 5B,C**). Taken together, these results show that in the absence of *Sox9, Nfib*, and *Nfix*, the ventral midbrain progenitors remained in the cell cycle at least until E15. However, by E18, progenitors were apparently in G0 in this region across all genotypes (**Fig. 3B**). This shows that the loss of *Nfib, Nfix* and *Sox9* postponed – but did not prevent – cell cycle exit in mD progenitors.

Nevertheless, this prolonged neurogenic window in the floor plate resulted in, on average, 60% more mDA neurons in E18 *Sox9NfibNfix*^*tcko*^ embryos (**Fig. 3C,D**). This increase was particularly prominent in the caudal VTA (**Fig. 3C**). As the mDA neurogenesis proceeds from rostral to caudal, with SNpc generated first and caudal VTA generated last (Bayer et al., 1995), we speculated that some of these excess mDA neurons in VTA could be generated after the normal mDA neurogenic window. To investigate this possibility, we administered BrdU in drinking water to pregnant females from E14-18, E15-18, and E16-18, and analysed the embryos for BrdU-labelled mDA neurons (**Fig. 3E**). As the number of mDA neurons had shown a modest but not statistically significant increase in *Sox9*^*cko*^ embryos compared to controls (**Fig. 3D**), we decided to focus only on *NfibNfix*^*dcko*^ and *Sox9NfibNfix*^*tcko*^ embryos. In the controls, we could find rare BrdU^+^Pitx3^+^TH^+^ neurons in the caudal VTA in the E14-E18 labelling scheme but none in later labelling windows, supporting the earlier findings that the majority of mDA neurons are generated by E14 (Bayer et al., 1995). In contrast, both *NfibNfix*^*dcko*^ and *Sox9NfibNfix*^*tcko*^ contained numerous BrdU-labelled mDA neurons, especially in E14-18 labelling, but also some in the E15-18 labelling window. These labelled cells showed highest enrichment in the caudal VTA. However, in the E16-18 labelling window no BrdU^+^ mDA neurons were present in either mutant. These results suggest that the loss of *Nfib/x* and *Sox9* prolongs the neurogenic window, contributing to the higher number of mDA neurons. Analysis of the more dorsal regions of these same samples showed that the *Sox9NfibNfix*^*tcko*^ dorsal midbrain generated neurons still in E16-18 stages (**Supp. Fig. 5D**), consistent with the 30-minute BrdU pulse labelling (**Fig. 2B**).

### *Sox9* loss leads to the appearance of ectopic mDA progenitors in late embryogenesis

One possible reason for the cessation of mDA neurogenesis in mutants is that *Sox9, Nfib*, and *Nfix* might have additional roles in postmitotic mDA differentiation, or that some environmental cues required for mDA neurogenesis are no longer present in later embryonic stages. Either one of these scenarios might result in mDA progenitors failing to mature into TH^+^ neurons.

Indeed, ectopic Sox2^+^ cells appeared below the floor plate VZ in *Sox9*^*cko*^ but particularly in *Sox9NfibNfix*^*tcko*^ mutants (**Supp. Fig. 6A**) from E14 onwards. In contrast, in *NfibNfix*^*dcko*^ mutants we could not detect a similarly drastic phenotype.

These ectopic cells might result from a structural collapse of the apical junctions in the most ventral VZ. At E13, before the appearance of these ectopic progenitors, adherens junction -marker beta-catenin remained normally expressed in the *Sox9NfibNfix*^*tcko*^ midbrain (**Supp. Fig. 6B**). Two days later, when ectopic progenitors could be seen in both *Sox9*^*cko*^ and in *Sox9NfibNfix*^*tcko*^, beta-catenin was lost specifically in *NfibNfix*^*dko*^ and in *Sox9NfibNfix*^*tcko*^ but remained surprisingly unaffected in *Sox9*^*cko*^ (**Supp. Fig. 6C**). Moreover, tight junction marker ZO1 was normally expressed at E15 in *Sox9NfibNfix*^*tcko*^ VZ (**Supp. Fig. 6D**). These results suggest that the loss of cell-to-cell adhesion may contribute to but cannot be the sole cause of these ectopic cells.

The ectopic cells lacked Pitx3 and TH but retained expression of mDA progenitor markers GFAP, Sox6, and FoxA2, and did not take up BrdU (**Supp. Fig. 6E**). This suggests that these cells could be postmitotic mDA progenitors unable to complete their maturation into mDA neurons.

### Ectopic Sox2^+^ cells in the dorsal adult *NfibNfix*^*dcko*^ aqueduct

As the inactivation of *Sox9* by En1^Cre^ results in death immediately after birth (Vong et al., 2015), we could not analyse the adult brain phenotype of *Sox9*^*cko*^ or *Sox9NfibNfix*^*tcko*^ mutants. However, *NfibNfix*^*dcko*^ mutants do survive, although most die before weaning. A previous study described a lack of foliation in the E18 *Nfib*^*-/-*^ cerebellum (Steele-Perkins et al., 2005). Consistent with this, the En1^Cre^-mediated conditional loss of *Nfib* led to a major cerebellar malformation (**Supp. Fig. 7A**), resulting in severe movement control problems. We could also see more BrdU^+^ nuclei within the mutant cerebellum, suggesting a prolonged neurogenesis and/or gliogenesis in this region (**Supp. Fig. 7B**).

Some of the *NfibNfix*^*dcko*^ mutants survived to adulthood, enabling analysis of the aqueduct although long-term experiments were impossible. We noticed that while the midbrain ependymal cells in these mutants expressed S100b, FoxJ1, and Vimentin similar to controls, the region above the dorsal aqueduct contained a large cluster of Sox2^+^ cells which were likely postmitotic due to the lack of CyclinD1 and BrdU-uptake (**Fig. 4A**).

**Figure 4.**
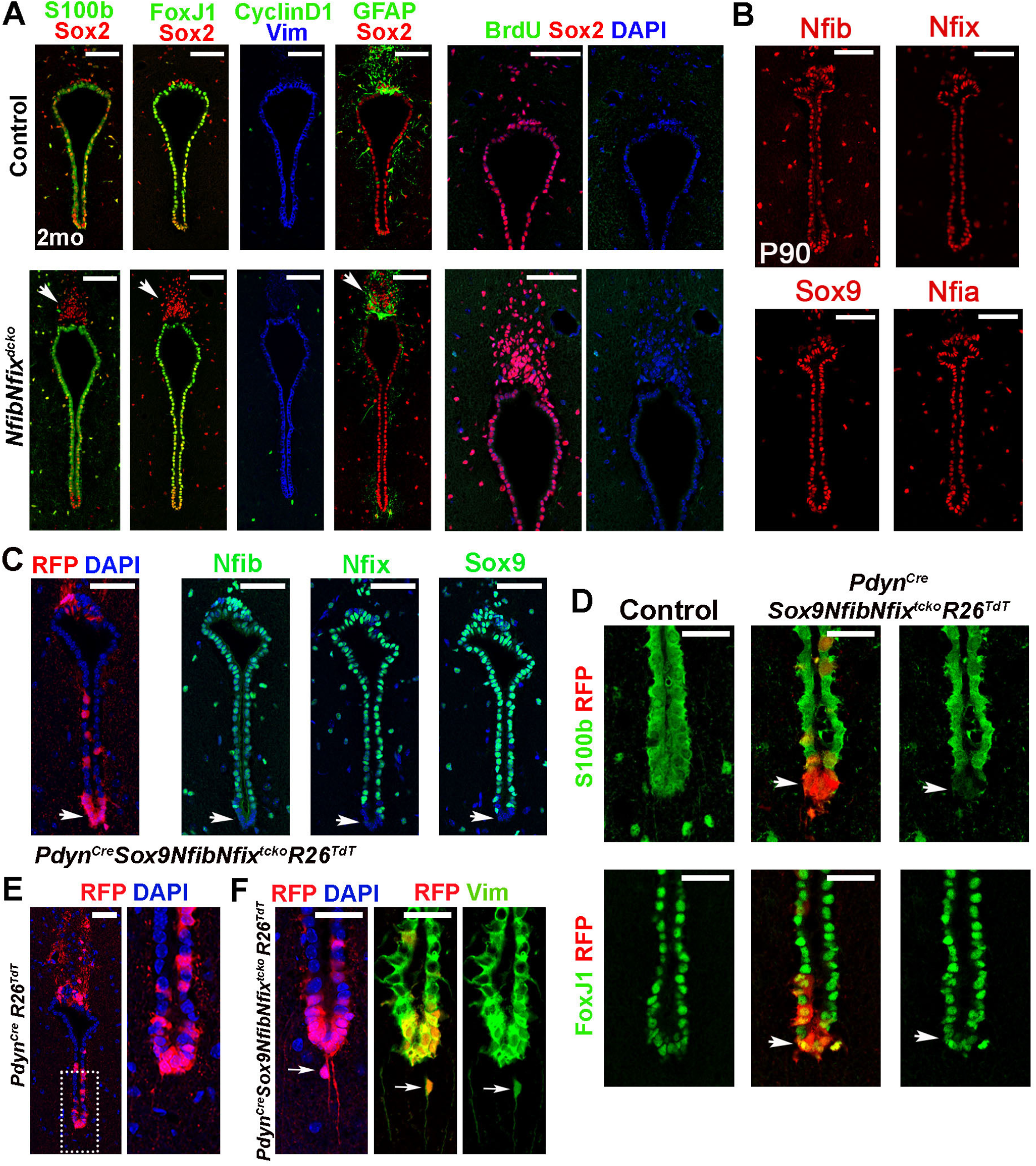
Nfib, Nfix and Sox9 are not required for the maintenance of G0 state in ependymal cells. **A)** IHC analysis of the aqueduct after 2 weeks of BrdU in drinking water. Arrows indicate the cluster Sox2^+^ cells above the aqueduct in mutants. **B)** IHC for Nfis and Sox9 in the adult aqueduct. **C)** Inactivation of *Nfib, Nfix* and *Sox9* flox alleles by Pdyn^Cre^ (arrows). **D)** IHC for ependymal markers S100b and FoxJ1 in control and *PdynCreSox9NfibNfix*^*tcko*^ aqueduct with arrows indicating the ventral-most region. **E)** Reporter expression in *Pdyn*^*Cre*^ *R26*^*TdT*^ aqueduct, with boxed area shown in high magnification. **F)** Ectopic nuclei (arrows) within the ventral RFP^+^ Vim^+^ fibers as occasionally seen below the *Pdyn*^*Cre*^*Sox9NfibNfix*^*tck*o^ aqueduct. Scalebars 50 μm in A-C and 25 μm in D-F.

### *Nfi* transcription factors or *Sox9* are not required for the maintenance of the G0 state

Next, we aimed to understand whether *Nfib, Nfix*, and *Sox9* not only induce but also maintain the G0 state in midbrain ependymal cells. Indeed, the expression of these factors was retained in the adult midbrain ependymal cells (Fig. 4B).

To inactive these genes in postmitotic ependymal precursors, we employed a *Pdyn*^*Cre*^ mouse line. *Pdyn*^*Cre*^ expression begins in the most ventral midbrain VZ at around E15-16 (**Supp. Fig. 7D**), a stage when almost no BrdU-uptake was detected in the control floorplate (**Fig. 3B**). In the adult, *Pdyn*^*Cre*^ label was seen in the most ventral and dorsal aqueduct in a *R26*^*TdT*^ reporter line (**Supp. Fig. 7D, Fig. 4C**). As expected, no mDA neurons were labelled with this Cre-line, demonstrating that *Pdyn*^*Cre*^ becomes active only after mDA progenitors had exited the cell cycle (**Supp. Fig. 7E**). The recombination of this Cre-line with floxed alleles of *Nfib, Nfix*, and *Sox9* lead to the absence of these factors in the mutant ventral aqueduct (**Fig. 4C**).

The lack of S100b but not FoxJ1 in the *Pdyn*^*Cre*^ *Sox-9NfibNfix*^*tcko*^ aqueduct indicated a maturation failure of the ependymal cells (**Fig. 4D**). Occasionally, ectopic nuclei within the ventral RFP^+^ Vimentin^+^ fibers were detected below the aqueduct (**Fig. 4E,F**). Although these fibers were found also in controls and will be described in greater detail further below, we could only detect the ectopic nuclei in *Pdyn*^*Cre*^ *Sox9NfibNfix*^*tcko*^ mutants. However, due to their rare appearance we could not determine their precise birth date by BrdU-pulse labelling.

### Dopamine signaling does not regulate the quiescent state of ependymal cells

Previous work suggested that similar to DA-feedback-regulated regeneration of mDA neurons in salamanders (Berg et al., 2011), evolutionary traces of this mechanism could still operate in the mammalian brain (Hedlund et al., 2016).

Intrigued by this idea, we hypothesized that if midbrain ependymal cells were not permanently in G0 but only quiescent, DA signaling might maintain the expression of quiescence-promoting transcription factors - such as *Nfis* and *Sox9*. Thus, in the absence of DA signaling, the expression of these factors would be downregulated. We thus generated local lesions of periaqueductal DA neurons by injecting 6-OHDA close to the aqueduct. Although this method successfully removed most TH^+^ cell bodies and fibers near the aqueduct, it had no discernable impact on the expression of *Nfis* or *Sox9* (**Supp. Fig. 7C**).

However, we still wanted to test whether the loss of DA signaling could influence ependymal cells in mice with inactivated *Nfib/x* and *Sox9*. For this, we repeated the 6-OHDA lesions in *Pdyn*^*Cre*^*Sox9NfibNfix*^*tcko*^ animals followed by BrdU in drinking water for 25 days to label any cells which might re-enter the cell cycle (**Supp. Fig. 7F**). However, we could not detect any increased BrdU labelling in any condition.

Taken together, these results indicate that while *Nfib, Nfix*, and *Sox9* are required for midbrain ependymal maturation, they are not essential for maintaining the G0 state.

### scRNAseq uncovers progenitor properties retained in a subset of adult midbrain ependymal cells

To better understand the properties of midbrain ependymal cells, we next analysed them by scRNAseq. FoxJ1^CreERT2^-labelled RFP^+^ ependymal cells were collected from lateral ventricles, 3rd ventricle, midbrain, and spinal cord, and analysed by SmartSeq2 (**Fig. 5A-C**). As an outgroup, cells from Nkx2.2^Cre^-labelled midbrains – mostly glia – were also included (**Fig. 5B,C**). After quality control 1473 cells, of which 567 were midbrain ependymal cells, were included in the analysis.

**Figure 5.**
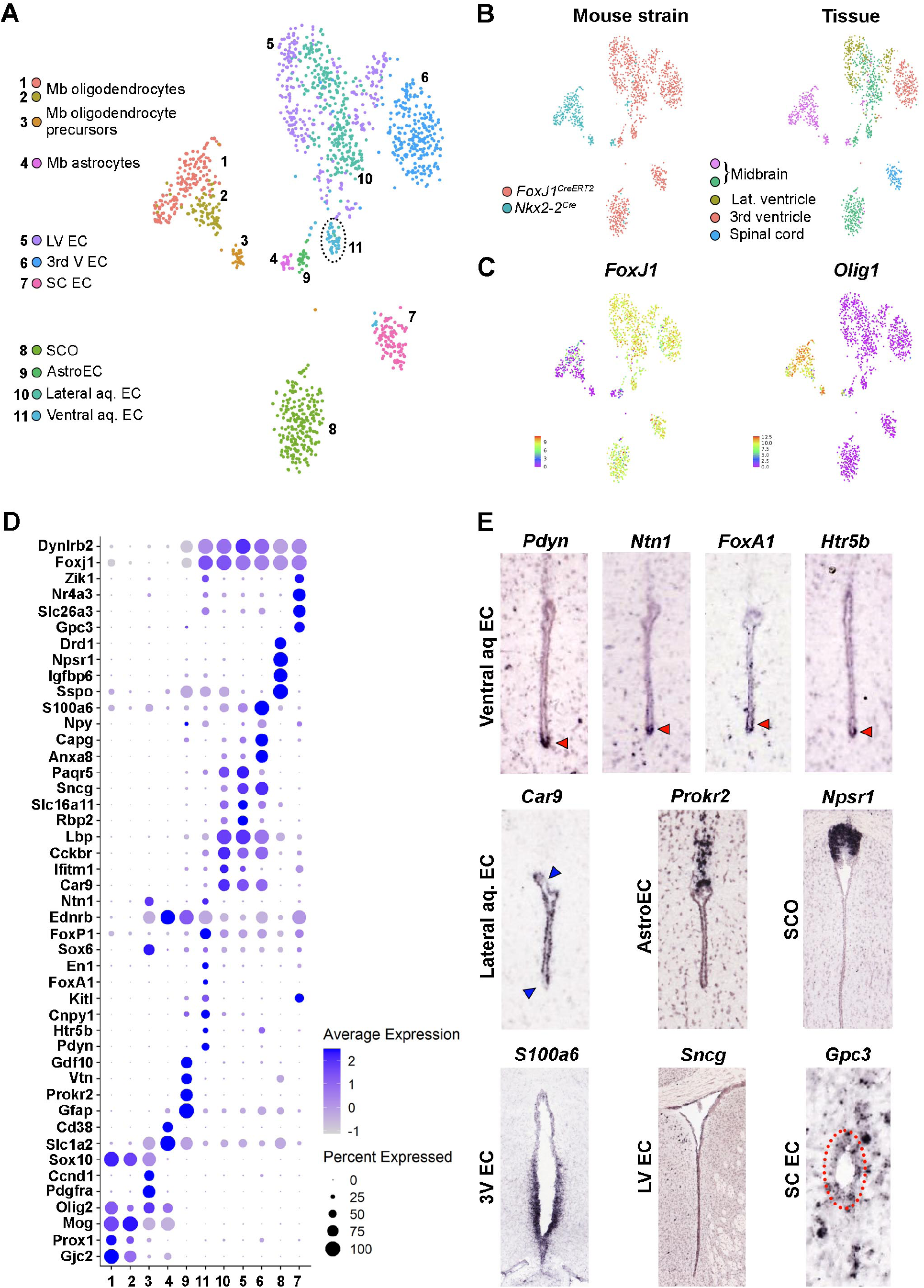
ScRNAseq analysis of the midbrain ependymal cells in the adult mouse. **A)** A tSNE plot of 1806 cells. Ependymal cells (EC) from the midbrain (Mb) aqueduct (Aq), lateral ventricles (LV), 3rd ventricle, and spinal cord (SC). SCO, subcommissural organ. The circle indicates the cluster corresponding to mDA-progenitor-derived ependymal cells. **B-C)** tSNE plots showing the mouse strains and tissue sources, and the two main cellular subtypes in the dataset. **D)** A dotplot of enriched genes in different clusters. **E)** Images from Allen Mouse Brain Atlas of ependymal cluster markers. Red arrowheads: the ventralmost ependymal cells; blue arrowheads: lower expression in the most ventral and dorsal tip; the red dotted circle: the spinal cord ependymal layer.

A tSNE projection segregated the dataset into 11 distinct clusters (**Fig. 5A**), each identifiable by its unique transcriptional profile, verified in situ in the Allen Mouse Brain Atlas (**Fig. 5D,E, Supp. Table 2**). The dataset can be explored by a web-based tool CellxGene (perlmannlab. org/resources).

In the tSNE projection the forebrain and lateral midbrain ependymal cells clustered together and shared the expression of several genes, while most dorsal and ventral ependymal cells formed three separate clusters (**Fig. 5A,D**). The identities of different clusters were annotated based on the expression of markers for distinct cell populations. Thus, cluster 8 expressing *Npsr1, Sspo* and *Igfbp6* corresponded to subcommissural organ (SCO), and the *Prokr2*^*+*^ *Vtn*^*+*^ cluster 9 consisted of astroependymal cells, located more caudal to SCO in the dorsal side (**Fig. 5A,D,E**). The former mDA progenitors in the ventralmost part of the aqueduct formed cluster 11 (**Fig. 5A**). Their enriched genes included opioid polypeptide hormone prodynorphin (*Pdyn*), a precursor of several endorphins; cytokine Kit ligand (*Kitl*, Stem cell factor); and serotonin receptor *Htr5b*, but also several genes and transcription factors typical of mDA progenitors, such as *Cnpy1, Ntn1, FoxP1, En1*, and *FoxA1* (**Fig. 5D,E**). This suggested that these cells might retain some mDA progenitor properties.

### Ventralmost ependymal cells retain several mDA progenitor markers in the adult

Our histological analysis revealed that the aqueduct was in close contact with GFAP^+^ astrocytes (**Fig. 6A,B**) and we could also detect Vim^+^ fibers protruding from the ependymal cells. These fibers were seen throughout the aqueduct but were particularly prominent in the dorsal and ventral tips. The dorsal fibers intermingled with GFAP^+^ fibers from the astroependymal cells (**Fig. 6A**). The dorsal and ventral ependymal cells also expressed Nestin, similar to the embryonic roof plate and floor plate, and the adult spinal cord, where Nestin^+^ fibers can be found in the dorsal side (Supp. Fig. 8A,B, Hamilton et al., 2009). The midbrain ependymal fibers mostly contacted nearby blood vessels, but on the ventral side, longer fibers contacted also PDGFR^+^ and Olig2^+^ cells - likely oligodendrocyte precursors (**Supp. Fig. 8D-F**).

**Figure 6.**
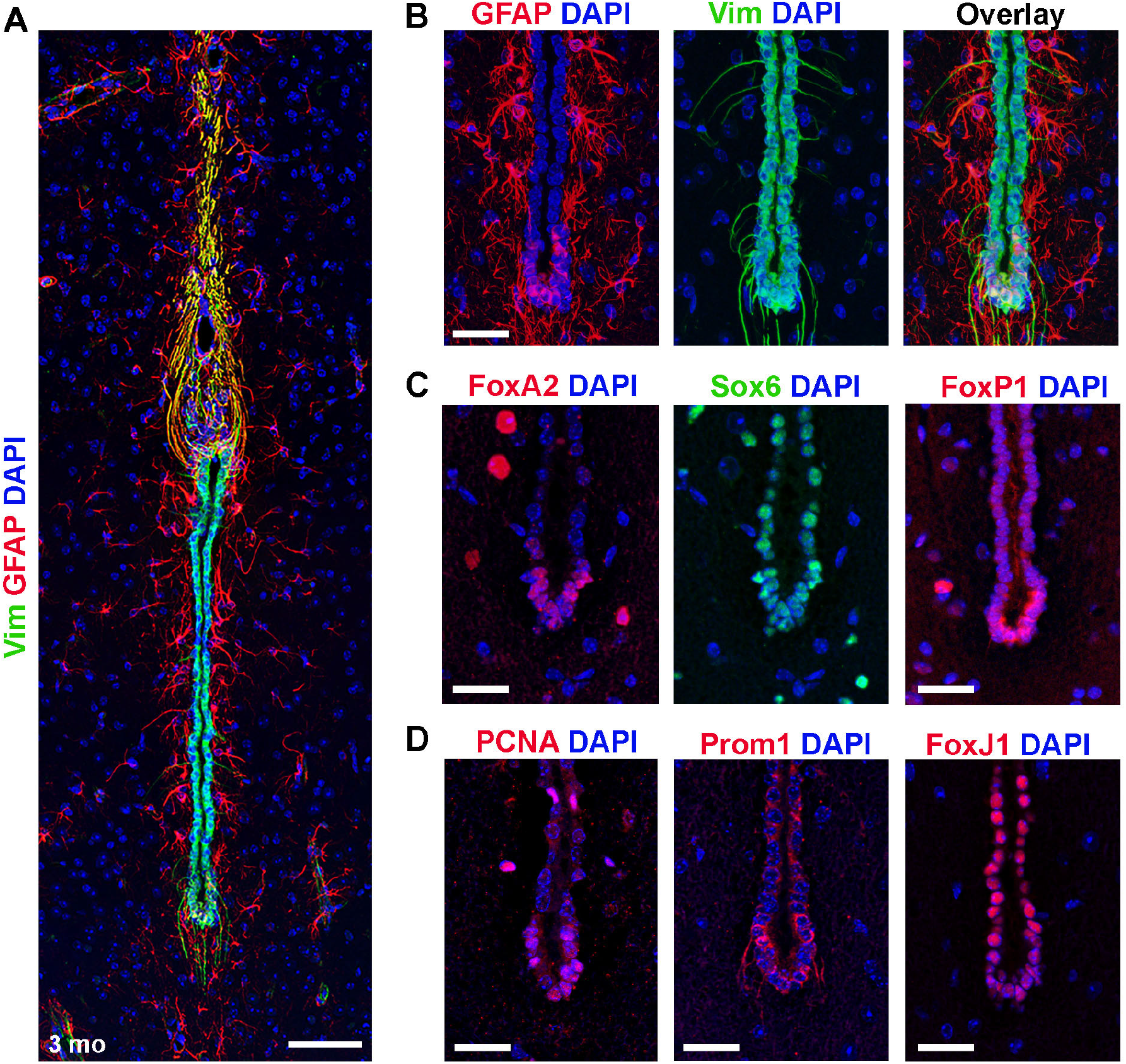
The ventral tip of aqueduct expresses mDA progenitor markers. **A)** An IHC staining of adult aqueduct showing long dorsal and ventral fibers extending from the ependymal cells, and astrocytes in close contact with the aqueduct. Maximum intensity projection. **B)** Close up of the ventral tip of the aqueduct in (A). **C-D)** IHC for mDA progenitor and neuronal progenitor markers and for ependymal marker FoxJ1. Scalebars 50 μm in A and 25 μm in B-D.

Supporting the results from scRNAseq, the ventralmost ependymal cells were FoxP1^+^ and Sox6^+^ (**Fig. 6C**). In addition to FoxA1, detected in the scRNAseq, the ventralmost cells were positive for mDA progenitor and neuron marker FoxA2 (**Fig. 6C**). They also displayed mosaic expression of progenitor markers GFAP, PCNA, and Prom1 (CD133), whereas ependymal marker FoxJ1 appeared weaker compared to more dorsal aqueduct (**Fig. 6B,D**). Mosaic expression of several markers suggests internal heterogeneity within this domain, which may either be temporal or represent subtypes of ependymal cells, perhaps with different properties.

### Midbrain ependymal cells are quiescent *in vivo* but can proliferate *in vitro*

Some spinal cord ependymal cells can slowly proliferate *in vivo*, as well as differentiate into both glia and neurons *in vitro*, and contribute to glial scar formation after a spinal cord injury (Meletis et al., 2008, Barnabe-Heider et al., 2010).

As we had seen several progenitor markers being expressed in the ventralmost domain of the aqueduct, we investigated whether these cells had any proliferative capacity *in vivo*. For this, we gave tamoxifen-treated *FoxJ1*^*CreERT2*^*R26*^*TdT*^ mice BrdU in drinking water for 6 weeks. Although this treatment successfully labelled both SVZ neural stem cells and spinal cord ependymal cells, BrdU uptake was not seen in any midbrain ependymal cells (**Supp. Fig. 8C**). These results suggest that although midbrain ependymal cells share several features with the ependymal cells in the spinal cord, they cannot proliferate *in vivo*, at least not under normal conditions. We speculated that some signals from the immediate surroundings might maintain the G0 state of these cells, and that removing them from this environment might trigger proliferation.

A previous study reported sphere-forming potential in different parts of the neuraxis, although those spheres were not genetically fate-mapped to verify their origin (Golmohammadi et al., 2008). To trace the midbrain ependymal cells *in vitro* we collected both the ependymal cells and the surrounding parenchymal cells from tamoxifen-treated *FoxJ1*^*CreERT2*^*R26*^*TdT*^ reporter mice (**Fig. 7A**). Cells were then plated both on fibronectin-treated surfaces and in free-floating cultures in neurosphere-media. Bipolar RFP^+^ cells were present on the fibronectin-coated plates in cultures from the aqueduct, but not from the parenchyma (**Fig. 7B**).

**Figure 7.**
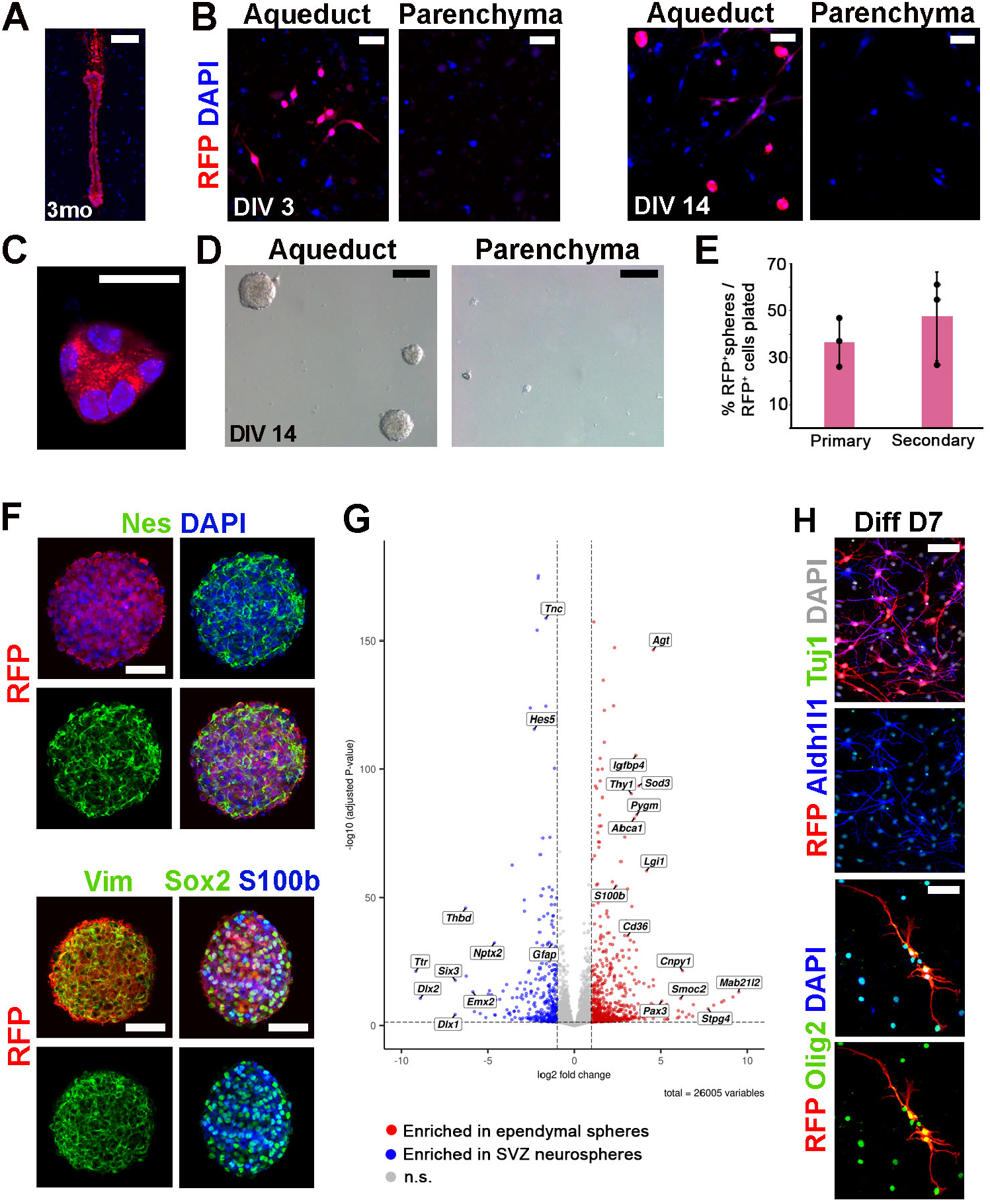
Midbrain ependymal cells proliferate and produce glial cells *in vitro*. **A)** RFP staining of the aqueduct of a tamoxifen-treated *FoxJ1*^*CreERT2*^ *R26*^*TdT*^ mouse. **B)** Aqueductal and parenchymal cells from the *FoxJ1*^*CreERT2*^ *R26*^*TdT*^ midbrain on fibronectin. **C)** An ependymal sphere after 2 weeks *in vitr*o on fibronectin. **D)** Ependymal and parenchymal free-floating cultures after 2 weeks. Quantification of RFP^+^ spheres from primary cultures on fibronectin (n=3 wells) and from secondary cultures on 96-well plates (n=3 plates). IHC on RFP^+^ spheres after 2 passages *in vitro*. **G)** A volcano-plot showing genes enriched in midbrain ependymal spheres vs SVZ -derived neurospheres from the same animals. **H)** Differentiated ependymal sphere cultures after 7 days. Scalebars 50 μm in A,F,H, 25 μm in B,C and 100 μm in D.

After 2 weeks *in vitro*, most of the parenchymal culture cells had died without forming any spheres in either fibronectin-coated wells or the free-floating cultures (**Fig. 7B,D**). In contrast, the aqueductal cultures formed small RFP^+^ spheres on fibronectin and larger spheres in free-floating cultures (**Fig. 7B-E**). They could also form secondary spheres (**Fig. 7E**) and could be passaged for several months.

The RFP^+^ spheres expressed several ependymal and neural progenitor markers, such as Nestin, Vimentin, Sox2, and S100beta (**Fig. 7F**). We compared their expression profiles by RNA sequencing to those of true SVZ-neurospheres from the same animals. Although spheres in a free-floating culture are always a mix of several cell types, we could see a clear transcriptional difference between the SVZ neurospheres and midbrain ependymal spheres (**Fig. 7G, Supp. table 3**). For example, SVZ neurospheres expressed more neural stem cell marker GFAP, and were enriched in forebrain-specific genes such as *Six3, Emx2, Dlx1* and *Dlx2*. The midbrain spheres in turn contained several ependymal markers typical for midbrain such as *Mab21l2, Pax3* and *Cnpy1*. By comparing these results to those from scRNAseq, we could see that spheres contained genes specific for all four different midbrain ependymal types (**Supp. Table 2**), indicating that ependymal cells in all different domains of the aqueduct could contribute to the sphere formation.

However, although ependymal spheres showed extensive proliferative capacity, they could only produce glial cells, not neurons, in the differentiating conditions (**Fig. 7H**).

Taken together, these results indicate that midbrain ependymal cells retain a capacity for proliferation if placed in a suitable environment, although their differentiation capacity *in vitro* is limited.

## Discussion

Several studies have highlighted the diverse roles of *Sox9* and *Nfi* transcription factors in embryonic neurogenesis and gliogenesis as well as in the maintenance of adult stem cell niche (Das Neves et al., 1999; Steele-Perkins et al., 2005; Campbell et al., 2008; Martynoga et al., 2013; reviewed in Harris et al., 2015; Sarkar and Hochedlinger 2013).

In the current work, we focus on the function of these factors in midbrain development and show that they are required to induce mitotic exit in the midbrain neuronal progenitors, and that their loss in the embryonic midbrain results in prolonged and increased neurogenesis, impaired gliogenesis, and delayed ependymal maturation. We also show that ependymal cells derived from mDA progenitors have radial glia-like characteristics, contact nearby blood vessels, and express a precursor for endorphins, suggesting that these cells might form a novel circumventricular organ in the adult.

### Nfib, Nfix, and Sox9 regulate the mitotic exit and ependymal maturation in the midbrain

Earlier studies on Sox9 and Nfi transcription factors in the developing CNS have focused on cortex, cerebellum, and spinal cord (Harris et al., 2015; Jo et al., 2014; Vong et al., 2015). Here we focused on the embryonic midbrain, and identified these factors as being upregulated in late versus early stage mDA progenitors.

Based on several of our observations, we postulate that these factors are required to initiate the mitotic exit and promote ependymal maturation. Firstly, they were gradually upregulated in the mDA progenitors during the neurogenic window and peaked as proliferation and neurogenesis ceased. Secondly, their conditional inactivation in the midbrain led to prolonged and increased proliferation and neurogenesis, and reduced expression of marker genes for mature ependymal cells. This effect was not restricted to the mDA domain but was seen throughout the midbrain.

This phenotype could be detected in single *Sox9*^*cko*^ mutants but required the inactivation of both *Nfib* and *Nfix*, suggesting a functional redundancy between Nfi transcription factors similar to the developing hippocampus (Harris et al., 2016). The function of *Nfia* in the midbrain development remains to be investigated.

Compared to earlier results from other brain regions, our results from the midbrain show both similarities and differences. Our data from *Sox9*^*cko*^ midbrain resembles the observations from the cerebellum in these same mutants (Vong et al., 2015), suggesting that Sox9 regulates mitotic exit and neurogenesis in both of these neighbouring tissues. In contrast, in the developing cortex, Sox9 is needed for the induction and maintenance of neural stem cells, and its loss results in reduced neurogenesis (Scott et al., 2010).

Nfi transcription factors, in turn, promote differentiation and suppress the self-renewal of neural progenitors in the embryonic forebrain (Piper et al., 2010, 2014; Heng et al., 2014; Harris et al., 2016). In *Nfix*^*-/-*^ and *Nfib*^*-/-*^ embryos, the number of telencephalic radial glial progenitors increases, but their maturation into intermediary progenitors is delayed, thus postponing and prolonging neurogenesis and contributing to brain malformations (Harris et al., 2016). In contrast, in the midbrain of *NfibNfix*^*dcko*^ mutants, the increased BrdU-uptake seen in the VZ did not delay or inhibit neurogenesis but led directly to increased production of neurons, resembling the phenotype in the *Sox9*^*cko*^ midbrain.

This increased neurogenesis may be further exacerbated in the lateral midbrain by drastically reduced gliogenesis in the mutants. If Nfib/x and Sox9 are already required in the progenitor cells to induce glial generation, their absence might switch gliogenic cell divisions to become neurogenic. Inactivation of these factors specifically in the postmitotic glial precursors would clarify the role of *Sox9* and *Nfis* in midbrain gliogenesis.

Although in *Sox9*^*cko*^ mutants we observed increased neurogenesis in the dorsolateral midbrain and prolonged BrdU-uptake in mDA progenitors, the increase mDA number by E18 in these mutants was relatively mild. In contrast, the loss *Nfib, Nfix*, and *Sox9* together resulted in a more severe mDA phenotype compared to *NfibNfix*^*dcko*^, indicating that Sox9 does contribute to the regulation of mDA neurogenesis. Furthermore, the inactivation of *Sox9* by itself or combined with *Nfib* and *Nfix* led to the accumulation of ectopic mDA progenitors after E14. Understanding the differences between these phenotypes and the transcriptional targets of *Nfis* and *Sox9* in mDA neurogenesis would benefit from scRNAseq-characterization of these mutants.

In the cortex, *Nfix* is required for the differentiation of ependymal cells and the maintenance of the adult ependymal layer (Harkins et al., 2022). Our data from the embryonic midbrain also show reduced expression of ependymal marker genes FoxJ1 and S100beta in the absence of *Nfib, Nfix*, and *Sox9*. However, as we could see a normal expression of these factors and an intact ependymal layer in the adult *NfibNfix*^*dcko*^ aqueduct, the ependymal maturation in these mutants was only postponed, likely due to the delayed mitotic exit. In contrast, Pdyn^Cre^-mediated inactivation of all three factors during late embryogenesis resulted in the loss of S100b. This indicates that Nfib, Nfix, and Sox9 promote mitotic exit enabling the ventricular cells to start their differentiation into ependymal cells. In addition, Sox9 appears to have an additional role in activating ependymal gene expression independently of FoxJ1, at least in the ventralmost aqueduct.

### Unique characteristics of ventralmost ependymal cells in the midbrain

Spinal cord ependymal cells have the capacity to proliferate and form both glia and neurons *in vitro*, and they contribute to the formation of glial scars after injury *in vivo* (Horner et al., 2000, Meletis et al., 2008, Ren et al., 2017). In contrast, both histological and scRNA sequencing analyses indicate that ependymal cells in the lateral ventricles neither proliferate nor contribute to neurogenesis or gliogenesis *in vivo* (Shah et al., 2018, Spassky et al., 2005). The properties of midbrain ependymal cells have not been characterised before in detail.

In this study, scRNAseq identified four distinct types of cells within the midbrain ependyma. While ependymal cells of the lateral walls of the aqueduct resembled transcriptionally the “classic” ependymal cells in the lateral ventricles, the dorsal and ventral tips of the aqueduct consisted of cells that shared many features with neural progenitor cells, such as the expression of Nestin and GFAP. The two dorsal cell types were astroependymoglial cells and the subcommissural organ (SCO), whereas the ventralmost cell population consisted of cells that descended from mDA progenitors.

The ventralmost *Lmx1a*^*+*^ ependymal cells characterized here also expressed *Gfap, Pcna, Prom1*, and several mDA progenitor markers such as *FoxA1/2, En1, Cnpy1, Sox6*, and *FoxP1*. As the expression of these factors was not uniform in this domain, there is likely further heterogeneity – possibly temporal – to be uncovered within this cluster. Despite their radial glia-like characteristics, these cells did not incorporate BrdU *in vivo*, nor did they express *CyclinD1*.

As these cells did form gliogenic spheres *in vitro*, they seem to reside in a state of deep quiescence, instead of being terminally in G0. This state might be maintained either by inhibitory signals from the environment or by the lack of cell cycle-promoting stimuli, or both. For example, inhibition could be based signals from SCO (Vera et al., 2013), or on contacts with the surrounding parenchyma and/or each other, as in a mouse model of hydrocephalus where the ependymal cells start to divide in order to repair the damaged ventricular walls (Batiz et al 2011).

In the adult salamander, the midbrain ependymoglial cells are kept in a quiescent state by feedback inhibition from the surrounding mDA neurons (Berg et al., 2011). Previous work (Hedlund et al., 2016) suggested that this mechanism might also generate some proliferating cells in the mouse midbrain. However, we noted no ependymal cell re-entry into the cell cycle after local DA depletion. Neither did *NfibNfix*^*dcko*^ nor *Pdyn*^*Cre*^*Sox9NfibNfix*^*tcko*^ adult mutants show any signs of cell cycle re-entry in the midbrain ependyma, suggesting that for the maintenance of G0 in the adult these factors are dispensable. Other mechanisms to keep these cells in a quiescent state might involve chromatin modifications or a switch of transcription factor complexes from the ones promoting initiation of G0 to the ones maintaining it.

Whether growth factor infusions or other stimuli can trigger the midbrain ependymal cells to re-enter the cell cycle remains to be investigated and should be accompanied by scRNAseq and a reporter line such as Fucci, or m-Venus-p27K (Oki et al., 2014) to track any changes in the cell status.

### Ventralmost aqueduct cells: a novel type of circumventricular organ?

In addition to radial glia and mDA progenitor marker expression, the ventralmost aqueductal cells contained long Vimentin^+^ fibers contacting blood vessels and glial cells, resembling tanycyte-like cells in several circumventricular organs (CVOs). CVOs, one of which is SCO, regulate various behaviors and body homeostasis (Benarroch 2011). CVOs can send and receive various signals, such as peptides and hormones, between peripheral blood and the central nervous system, thus bypassing the blood-brain barrier. Some contain elongated, Nestin^+^ GFAP^+^ tanycyte-like cells, which can be triggered by mitogens to incorporate BrdU (Furube et al., 2020). In addition, sphere cultures have been obtained from different anteroposterior levels of the neuraxis, although they have not been fate-mapped to any CVOs (Weiss et al., 1996, Golhohammadi et al., 2008). Taken together, some cells within CVOs possess multiple radial glia or tanycyte-like properties and a potential to cell cycle re-entry, suggesting that they are not in permanent cell cycle arrest (Bennett et al., 2009).

Our results suggest that the ventralmost cells in the aqueduct might form a novel part of the CVO system. First, their radial glia/tanycyte-like morphology and gene expression profile resemble that of several CVOs. Second, in addition to their contacts with blood vessels and oligodendrocytes, the ventralmost aqueduct cells are surrounded by astrocytes, all of which could be potential signaling partners. Third, these cells express *Prodynorphin (Pdyn)*, a precursor of dynorphins, which are opioid neuropeptides involved in, for example, addiction, learning, stress and pain response, as well as mood regulation (Schwarzer 2009). Together these results indicate that these cells form a type of CVO that might respond to serotonin, or some other signaling, by secreting neuropeptides.

The main receptor for dynorphins is the kappa opioid receptor (KOR), encoded by *Oprk1*. KOR is highly enriched in the VTA mDA neurons, and dynorphin-KOR-signaling in VTA mediates the processing of aversive stimuli and promotes stress-induced compulsive behavior (Margolis and Karkhanis, 2019; Abraham et al., 2018). Dynorphin inputs to mDA neurons originate from various brain regions such as the striatum, lateral hypothalamus, amygdala, and the bed nucleus of stria terminalis (Margolis and Karkhanis, 2019). Although we do not yet understand the full extent and contacts of the fibers in ventral aqueduct cells, it is intriguing to speculate that some of these fibers might contact VTA DA neurons and possibly form a local input source of dynorphins to modulate their function.

Taken together, our findings provide critical insights into the regulatory networks governing midbrain neurogenesis and gliogenesis, with potential implications for understanding neurodevelopmental disorders and neurodegenerative conditions such as Parkinson’s disease. Understanding the details of mDA neuron development has been instrumental in developing protocols for engineering therapeutic mDA neurons from stem cells. Future studies will aim to explore how manipulating these pathways might offer therapeutic avenues by enabling more efficient stem cell protocols or by promoting endogenous brain repair.

## Materials and Methods

### Mouse lines and genotyping

*En1*^*Cre*^ (Kimmel et al., 2000, IMSR_JAX:007916), *Pdyn*^*Cre*^ (Krashes et al., 2014 IMSR_JAX:027958,) *Lmx1a*^*CreERT2*^ (Kee et al., 2017), *FoxJ1*^*CreERT2*^ (Meletis et al., 2008), *Nkx2-2*^*Cre*^ (Balderes et al., 2013), *Nfib flox* (Hsu et al., 2011), *Nfix flox* (Campbell et al., 2008), *Sox9 flox* (Akiyama et al., 2002, IMSR_JAX:013106), *R26*^*TdTomato*^ (Madisen et al., 2010, IMSR_JAX:007909; referred to as R26^TdT^ here), *R26*^*TrapCherry*^ (Hupe et al., 2014; referred to as R26^TrapC^ here) mouse strains and their genotyping have been described before. The mice were maintained in an outbred background. The noon of the day of the vaginal plug was considered as embryonic day (E) 0.5. Control animals or embryos were Cre-negative littermates. All experimental procedures followed the guidelines and recommendations of Swedish animal protection legislation and were approved by Stockholm North Animal Ethics board (permits 13830/18 and 16527-2023).

### Tissue processing of embryos and adult mouse brains

The embryos were collected in ice-cold DPBS (Gibco 14190144) and fixed in freshly prepared 4% paraformaldehyde (PFA, Sigma P6148) in DPBS in standard biopsy cassettes at room temperature (RT) over 1-2 nights depending on the embryonic stage, with PFA solution changed every day. The adult mice were deeply anaesthetized and intracardially perfused first +37°C DPBS followed by +37°C 4% PFA in DPBS, the brain dissected out and the entire midbain separated by coronal cuts using a brain matrix (AgnThos 69-2165-1) and postfixed over 2 nights at RT, changing PFA daily. The samples were dehydrated and processed into Histosec wax without DMSO (Merck 101676) using Leica automated tissue processor TP1020. The sections were cut at 5 μm (embryos) and 6-10 μm (adults) using automated rotating waterfall microtome (Epredia HM355S), collected on Superfrost Plus slides (Menzel-Gläser 631-9483), and dried overnight in a vertical position at +37°C. For the free-floating sections, the perfused and postfixed midbrains were cryoprotected in 30% sucrose, cut at 35 μm on a sliding microtome, and collected on a 48-well plate for staining.

### Immunohistochemistry and in situ hybridization

Fluorescent immunohistochemistry and in situ hybridization on sections were performed as described (Tiklova et al., 2019). Probes for *Dll1, Ascl1* and *Hes5* have been previously described (Saarimäki-Vire et al., 2007). Probe for *Neurog2* consisted of a synthetized 808 bp GeneArt Strings (Invitrogen) fragment of mouse *Neurog2* cDNA, from CCCTTCTCCACCTTCCTCCT to ATGCCTATTGTC-CCGCCCTT. The fragment was cloned into a pCR-Blunt II TOPO vector (Invitrogen K280002) according to manufacturer’s instructions.

The free-floating adult brain sections were permeabilized in 0.3% Triton X-100 in PBS for 10 mins, blocked in 5% donkey serum (Jackson Immunoresearch 017-000-121) in 0.1% TX-100 in DPBS for 1 hour at RT and incubated in primary antibody in blocking solution o/n at RT, washed several times in 0.1% Triton X-100 in PBS followed by incubation in the biotinylated secondary antibody and then stained using DAB-detection kit (Vector SK-4100) according to manufacturer’s instructions.

For immunocytochemistry, the cells were fixed 30 minutes RT in 4% PFA, washed several times in PBS, permeabilized in 0.3% Triton X-100 for 10 minutes and blocked in 5% donkey serum in 0.1% TX-100 in DPBS for 1 hour at RT. The samples were incubated in primary antibodies o/n +4C, washed several times in 0.1% Triton X-100 in PBS, and incubated in secondary antibodies for 2 hours RT. The whole ependymal spheres from free-floating cultures were fixed, dehydrated and processed into paraffin manually, and then sectioned and stained following the protocol for embryonic and adult brain samples.

The information about primary and secondary antibodies is found in Supp. table 4.

### Cell culture

The adult mice were euthanized by CO2, decapitated, and brain tissue collected in ice-cold DPBS. The brains were cut coronally at 1 mm using a pre-cooled brain matrix (AgnThos 69-2165-1) and razor blades. The aqueduct and parenchymal tissue from the midbrain and subventricular zone from the lateral ventricles were microdissected from the sections using sterile 27G needles attached to 1 ml syringes.

For primary cultures of ependymal and parenchymal cells on fibronectin, the tissues were enzymatically dissociated using a papain kit (Miltenyi 130-092-628) with manufacturer’s instructions, followed by myelin-removal using beads (Miltenyi 130-096-731) and cells plated on fibronectin-coated 96-well plates at 20 000 cells / well in NeuroCult Proliferation solution (Stem Cell Technologies 05702) with heparin (Stem Cell Technologies 07980), bFGF (Stem Cell Technologies 78003), EGF (Stem Cell Technologies 78006.1) according to manufacturer’s instructions, with antibiotic-antimycotic solution (Gibco 15240096).

For free-floating sphere cultures, the tissues were dissociated into single cell suspension using NeuroCult enzymatic dissociation kit for adult mouse and rat CNS tissue (Stem Cell Technologies 05715) and plated on ultra-low attachment 6-well plates (Corning 3471) in NeuroCult Proliferation solution with growth factors and antibiotic-anti-mycotic as described above. The spheres were split using either same dissociation kit as above, or DPBS with 0.02% EDTA for 15-20 minutes RT followed by trituration (Sigma E8008). The ependymal spheres were split at 5-7 days interval, the neurospheres from SVZ 3-5 days interval depending on the sphere size and appearance. For secondary spheres formation test the single cell suspension was plated on round bottomed 96-well plates on a clonal density, or cells were FAC sorted for RFP directly on a 96-well plate. The plates were monitored after the cells had settled to verify the absence of any spheres at this stage. The plates were analysed 10 days later and only spheres larger than 50 μm in diameter were counted.

Spheres were collected for RNA sequencing from both neurosphere and ependymal sphere cultures from the first passage, and were snapfrozen and kept in -80C until RNA extraction.

For differentiation, the dissociated spheres were plated on 8-chamber slides (Millipore PEZGS0816) coated on a thin layer of Matrigel without growth factors (Corning CLS356230). One day after plating the media was changed to NeuroCult differentiation solution (Stem Cell Technologies 05704) with antibiotic-antimycotic.

### Laser-capture microdissection

Tissue for laser capture microdissection was collected from fresh-frozen embryos (E11 n=4, E13 n=5, E18 n=5) which were embedded in OCT, cut coronally at 10 μm, and collected on PEN membrane-coated slides (Zeiss 415190-9042-000), which were pre-treated under UV-light. The sections were fixed in cold (−20°C) 95% EtOH, OCT rinsed off in PBS for 2 mins, and sections dehydrated in 70%, 95% and absolute EtOH for 30 seconds each and quickly air-dried. The mDA progenitor domain was identified by direct fluorescence and captured from several sections across the midbrain using Leica LMD7000 system using 20x magnification and keeping the laser power as low as possible. The tissue pieces were collected in 0.2 ml low-adhesive tubes (VWR 732-0548). Lysis buffer (0.4% Triton X-100 with 2U/μl RNAse inhibitor, Takara) was added immediately after sample tube removal from the machine, the tubes spun down briefly and snap-frozen on dry ice. The samples were stored at -80°C until library preparation.

### Isolation of cells for scRNAseq

Ependymal and glial cells for scRNA sequencing were prepared from freshly dissected brains. Ependymal tissue was microdissected from the lateral ventricles, midbrain aqueduct, third ventricle, and spinal cord from 3-month-old *FoxJ1*^*CreERT2*^ *R26*^*TdT*^ animals (n=10; 5 females & 5 males). Midbrain tissue was also microdissected from 3-month-old *Nkx2-2*^*Cre*^ *R26*^*TrapC*^ mice (n=7, all females). Tissues were dissociated using papain kit (Miltenyi 130-092-628 for ependymal cells, Worthington LK003150 for *Nkx2-2*^*Cre*^ midbrains). FACS buffer was HBSS (Gibco) without Ca^2+^ and Mg^2+^, with 50 mM glucose added to support ependymal cells. Viability dye TO-PRO3 Ready Flow Reagent (Invitrogen R37170) was added 15 minutes before sorting. Cells were sorted using BD FACS Aria III (BD Biosciences) and collected on 96-well PCR plates. The nozzle size was 100 for *FoxJ1*^*CreERT2*^ samples and 130 for the *Nkx2-2*^*Cre*^ samples.

### RNA sequencing and bioinformatic analyses

The RNA was extracted and cDNA libraries prepared using standard SmartSeq2 protocol using Tn5 made in the lab (Picelli et al., 2013, 2014). The quality of cDNA and tagmented cDNA was checked on a High-Sensitivity DNA chip (Agilent Bioanalyzer). LCM and scRNAseq samples were sequenced with Illumina HiSeq 2000, and spheres with Illumina NextSeq 2000.

DESeq2 package version 1.34.0 was used to analyze RNA-seq data, including differential gene expression, using a Negative Binomial GLM fitting per gene to account for overdispersion in the count data. Next, Wald statistical test was used to derive a p-value for each model coefficient and p-value adjustment for multiple testing correction with Benjamini & Hochberg (FDR) correction (Love et al., 2014). For neurospheres DE, lfcShrink() function in DESeq2 package was used to estimate the log fold changes (LFCs) more accurately and shrink the exaggerated LFCs which are mainly for genes with low counts or high variability. The adaptive t prior shrinkage estimator type=“apeglm” was used from the “apeglm” package (Zhu et al., 2018). The ‘EnhancedVolcano’ package version 1.10.0 was used to make volcano plots (Blighe et al., 2024).

For the analysis of single cell RNAseq, the sequenced reads were aligned to the mouse genome (mm10) that was merged with the eGFP sequence using the Star v2.3.1o (Dobin et al., 2013) and filtered for uniquely mapping reads. The expression values were then calculated for each Ensembl ID (release 69) based on the kilobase gene model with million mappable reads using ‘rpkmforgenes’ (Ramsköld et al., 2009). The low quality cells were then filtered based on the following thresholds: > 28.1% uniquely mapped reads, >49.5% mapped to exons, <12.6% reads mapped to the 3’ end, >3.2% of all genes detected and at least with a 100,000 normalisation reads. From a total of 1805 cells, 332 were removed with a resulting 1473 good quality cells that were used for the downstream analysis. Many of the QC metrics of the cells were monitored and checked as suggested by the ‘Scater’ R package (McCarthy et al., 2017). The gene Yam1 that codes for an lncRNA was removed from the dataset as the average expression of this gene in the cells compared to all the other genes was disproportionately high.

The clustering of the cells were done in two different steps: 1. The cells within each of the different tissue types were clustered individually, followed by 2: the cells from all the tissues were clustered together. The cells within each tissue type were clustered with the t-SNE + k-means algorithm. The differentially expressed genes for each of the different clusters in each tissue type were then detected using MAST (Finak et al., 2015). For the clusters in the midbrain tissue, differentially expressed genes were calculated similarly for each pairwise cluster comparisons. For clustering all the tissue types together, the shared nearest neighbor algorithm was applied with 5 neighbors to be considered for the graph construction. This graph was then used to cluster the cells with the ‘Louvain’ algorithm (Blondel et al., 2008). Based on the information from the tissue-specific clusters along with the information of the marker genes, 3 of the clusters one each from midbrain, forebrain and 3rd ventricle were merged with their respective closest cluster. Similar to the tissue-specific clusters, differential expression analysis was then carried out for each cluster using MAST. All of the above mentioned analysis and the visualizations were carried out through ‘Seurat’ R package v3 (Stuart et al., 2019)

The dot plot for scRNAseq data was made using Seurat package v4 (Hao et al., 2021).

The interactive single cell dataset exploration tool was created using CellxGene (https://github.com/chanzuckerberg/cellxgene).

### Microscopy and image analysis

The fluorescent images were taken on Zeiss confocal microscope LSM700, the whole ependymal spheres were imaged using Zeiss Observer Z.1, and the free-floating sections using Nikon Eclipse E1000. The images were processed, and brightness and contrast adjusted, either in Photoshop (Adobe) or in ImageJ2. If needed, an unsharp mask was also applied in Photoshop. Tiled images were compiled using either automated photomerge -function in Photoshop, or by pairwise stitching (Preibisch et al., 2009) in ImageJ2. Confocal stacks were compiled into maximum intensity projections using ImageJ2.

The quantification of BrdU uptake in ventricular cells (Figs. 2, 3 and Supp. Fig. 4) was analysed in ImageJ2 by measuring the proportion of BrdU^+^ signal to Sox2^+^ signal area. The regions of interests (ROIs) were first drawn manually following the outlines of Sox2^+^ VZ, divided into floor plate, roof plate, ventrolateral and dorsolateral domains. The signal was analysed in each embryo from several sections at 40 μm interval from anterior to posterior midbrain, and values of these sections averaged for each embryo for the statistical analyses.

The Olig2^+^ and Pou4f1^+^ cell numbers (Fig. 2) were counted as individual particles per section using Analyse Particles -function in ImageJ2 after automated thresholding. Three midbrain sections at 40 μm interval from each embryo were analysed.

DA neurons were counted as individual particles from each section (Fig. 3). Sections spanning the entire mDA domain in the antero-posterior direction were collected and analysed at 40 μm interval from each embryo (19-24 sections/embryo depending on the slight changes in the sectioning angle). First, ROI was drawn manually in ImageJ2 using the boundaries of TH signal for each section. Then Pitx3^+^ particle number was counted within this ROI. If Analyse Particles -function, even combined with Watershed function, was unable to reliably resolve areas where where DA neurons were densely clustered together, those sections were re-counted manually.

BrdU-labelling in DA neurons (Fig. 3) was counted manually in ImageJ2, checking each BrdU^+^ cell for signal in all channels, and again sections spanning the entire midbrain were analysed.

The statistical analyses were done in Minitab (Minitab. com) using default parameters for One-way ANOVA with Tukey simultaneous test for differences of means. First the data were verified to be homoscedastic and normally distributed using Levene and Ryan-Joiner (similar to Shapiro-Wilk) tests, respectively. The p-values from ANOVA and padj-values from Tukey are found in Supp. table 5 together with primary data from biological replicates.

### 6-OHDA lesioning, tamoxifen and BrdU treatment of mice

For lesioning of DA neurons in PAG, mice (n=2 for each genotype and condition, both males and females) were deeply anaesthetized with 2-2.5% isoflurane (KDG9623, Baxter Medical AB), their skin on top of the skull shaved and desinfected, and they were placed in a stereotaxic frame on a heating pad and given local anesthesia with s.c. injection of Marcain (169912, AstraZeneca). They were then unilaterally injected with either 2 μg (freebase amount) freshly prepared 6-OHDA (Sigma H4381) in 0.01% ascorbic acid (Sigma A92902) or the same volume of only 0.01% ascorbic acid at 0.1 μl/min. The injection coordinates were AP -3.1, ML 0.1, DV -3 relative to bregma (Paxinos and Franklin 2013). The needle was kept in place 1 minute before and 5 minutes after the injection. For relieving postoperative pain, the mice were given Rimadyl (14920, Zoetis) s.c. directly after the injection and 24 hours after the operation. PAG lesioning did not lead to any motor defects or other health issues.

Tamoxifen (Sigma T5648) was dissolved at 20 mg/ml in peanut oil (Sigma P2144) and incubated at +37C in a tube roller for 2-3 hours, protected from light, until it was completely dissolved, and the solution was used fresh. For the activation of Cre in the adult mice, *FoxJ1*^*CreERT2*^ *R26*^*TdT*^ animals received 3 mg tamoxifen in oral cavage on 3 separate days with a minimum of 2-day interval. For Lmx1a^CreERT2^ activation in the embryos, the pregnant females were treated with 3 mg tamoxifen in oral cavage when embryos were E9.5. BrdU (Roche 10280878001) was given either as an intraperitoneal injection in PBS (100mg/kg of body weight) or in drinking water (0.8mg/ml, with added sucrose to cover the taste). The BrdU water bottle was protected from light and water changed every 3 days.

## Acknowledgements

The authors thank Belinda Pannagel and Javier Avila-Cariño for the expert technical assistance with FACS; Juha Partanen for *Nkx2-2*^*Cre*^ mice and ISH probes; Jonas Frisén for *FoxJ1*^*CreERT2*^ mice; Jonas Muhr for the Sox6 antibody; Danijal Topcic for technical advice with neurosphere cultures; as well as both current and former members of Thomas Perlmann and András Simon labs for discussions and comments. The computations were performed on resources provided by SNIC through Uppsala Multi-disciplinary Center for Advanced Computational Science (UPPMAX) under Project SNIC 2017/7-348. The authors acknowledge support from Science for Life Laboratory, the National Genomics Infrastructure, NGI, and Uppmax for providing assistance in massive parallel sequencing and computational infrastructure. The authors thank especially Lokeshwaran Manoharan for the bioinformatic analyses of the ependymal data set.

## Competing interests

The authors declare no competing interests.

## Funding

The work was financed by Vetenskapsrådet (VR 2020-00884 T.P.; 2016-02506, T.P., L.L.), The Swedish Brain Foundation (Hjärnfonden, T.P.), Knut and Alice Wallenberg Foundation (T.P.), Torsten Söderbergs Foundation (T.P.), and Sigrid Juselius Foundation (Postdoctoral Fellowship, L.L.).

## Data availability

All RNA sequencing data will be made publicly available at the time of publication.

## Supplementary figures

**Supplementary Figure 1.**
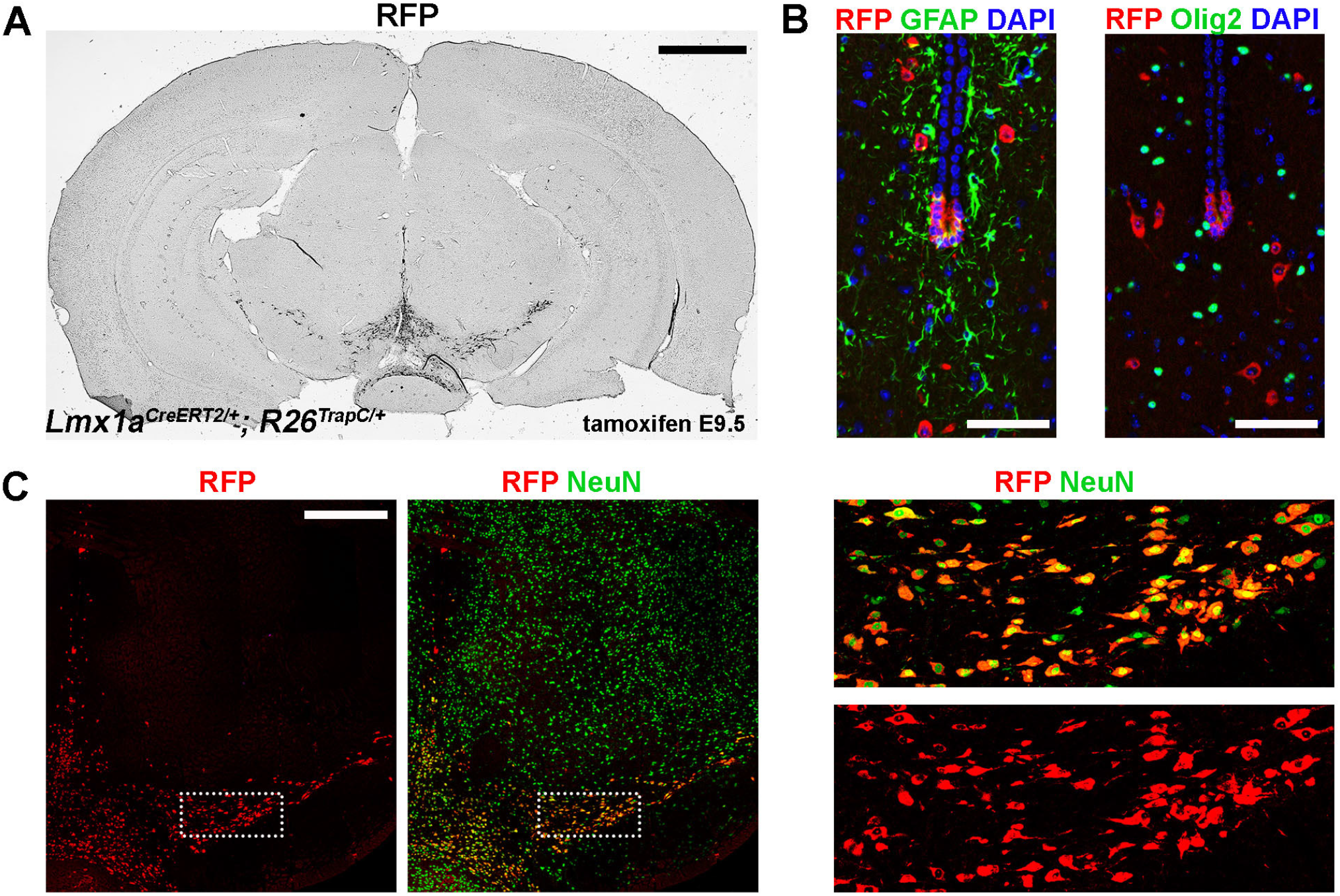
Dopaminergic progenitor domain does not generate glia. **A)** Chromogenic IHC staining of RFP on a free-floating section of an adult Lmx1a^CreERT2^ R26^Cherry^ reporter. **B)** Immunostaining of the ventral part of adult Lmx1a^CreERT2^ reporter aqueduct. GFAP expressing astrocytes or Olig2 expressing oligodendrocytes were not labelled with Cre. **C)** Ventral midbrain of Lmx1a^CreERT2^ reporter stained with antibodies against RFP and general neuronal marker NeuN. The boxed area is shown with high magnification on the right. Cells labelled with Lmx1a^CreERT2^ co-express RFP and NeuN, and had neuronal morphology, indicating a neuronal identity. In all samples, Cre was activated with a tamoxifen at E9.5. Scalebars 1 mm in A, 500 μm in C and 50 μm in B.

**Supplementary Figure 2.**
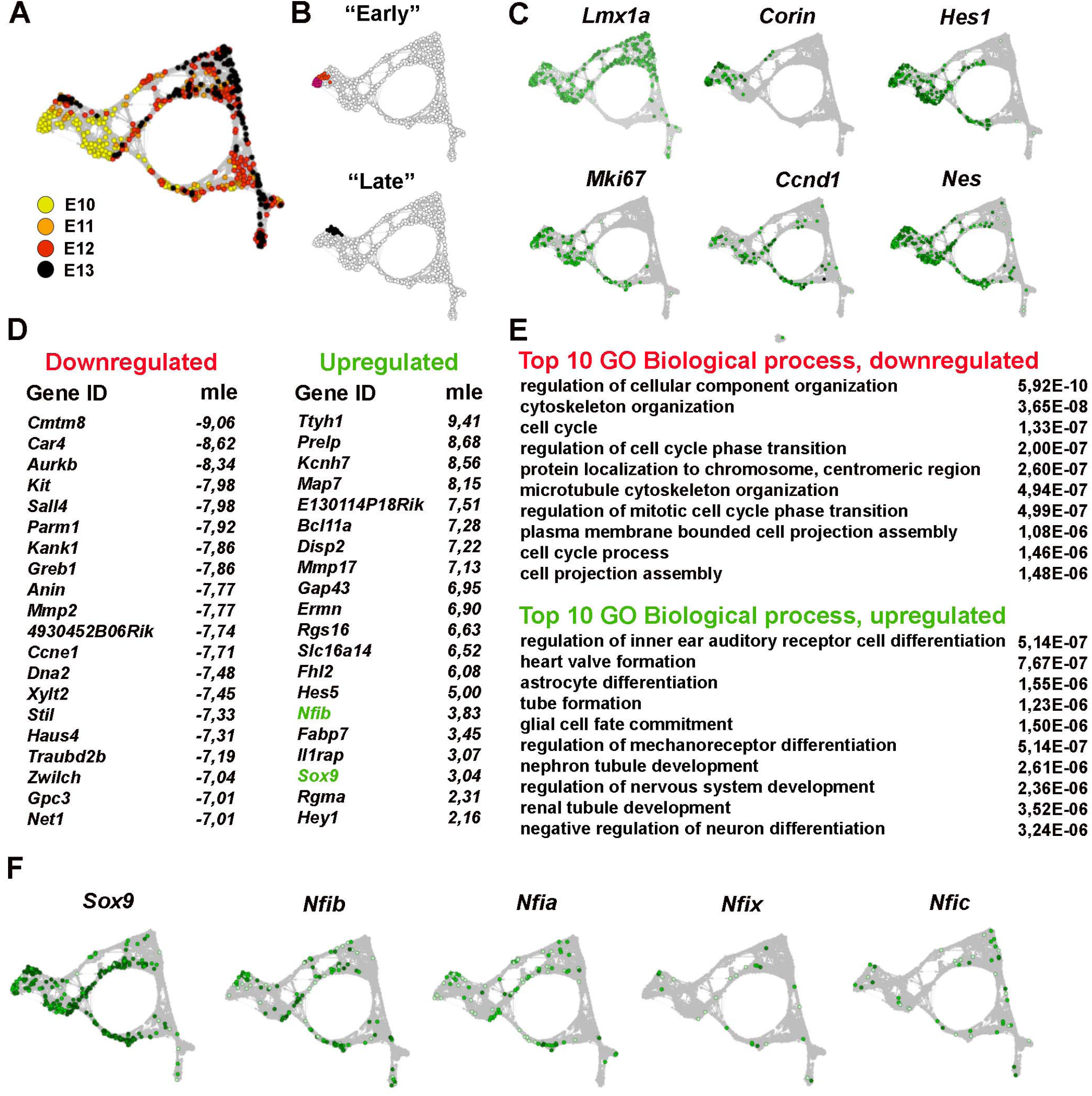
Upregulation of *Nfib* and *Sox9* in dopamine progenitors. **A)** A t-SNE plot from rShiny app from previously published single cell RNA sequencing data of early *Lmx1a*^*+*^ progenitors and precursors (Kee et al., 2017) showing the embryonic stages in the data set. **B)** Clusters used for the SCDE analysis in this study. The black cluster represents late stage *Lmx1a*^*+*^ progenitors, whereas the two red clusters were combined to represent early mDA progenitors. **C)** Genes expressed in proliferating mDA progenitors plotted in the rShiny app. **D)** Top 20 up- and down-regulated genes from SCDE analysis comparing early and late mDA progenitors. **E)** Top 10 GO-terms from Panther statistical overrepresentation test for Biological process run with default parameters. **F)** *Sox9* and different *Nfi* factors plotted on the rShiny app.

**Supplementary Figure 3.**
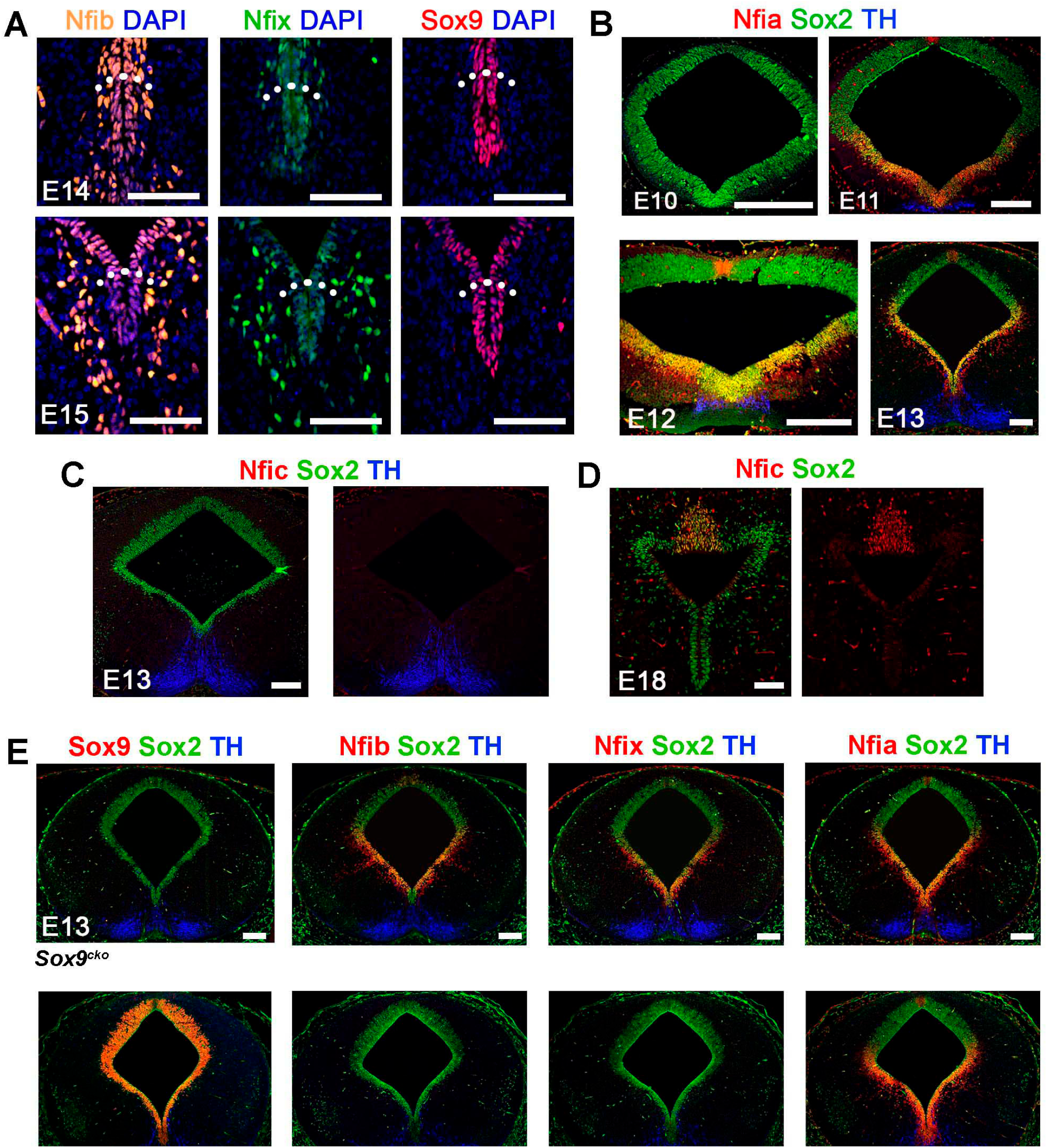
Expression analysis of Nfi and Sox9 transcription factors in the midbrain. **A)** IHC for Nfib, Nfix, and Sox9 in the quiescent DA progenitor domain, indicated by the dotted line. **B)** IHC analysis of Nfia in the early midbrain. **C-D)** IHC staining for Nfic showing no signal in the midbrain at E13 and expression in the most dorsalmost midbrain at E18. **E)** Expression of Sox9 and Nfi factors in different mutant embryos. Inactivation of *Sox9* does not affect the expression of *Nfi* factors, and *vice versa*. Scalebars 100 μm in A, 200 μm in B,C,E and 50 μm in D.

**Supplementary Figure 4.**
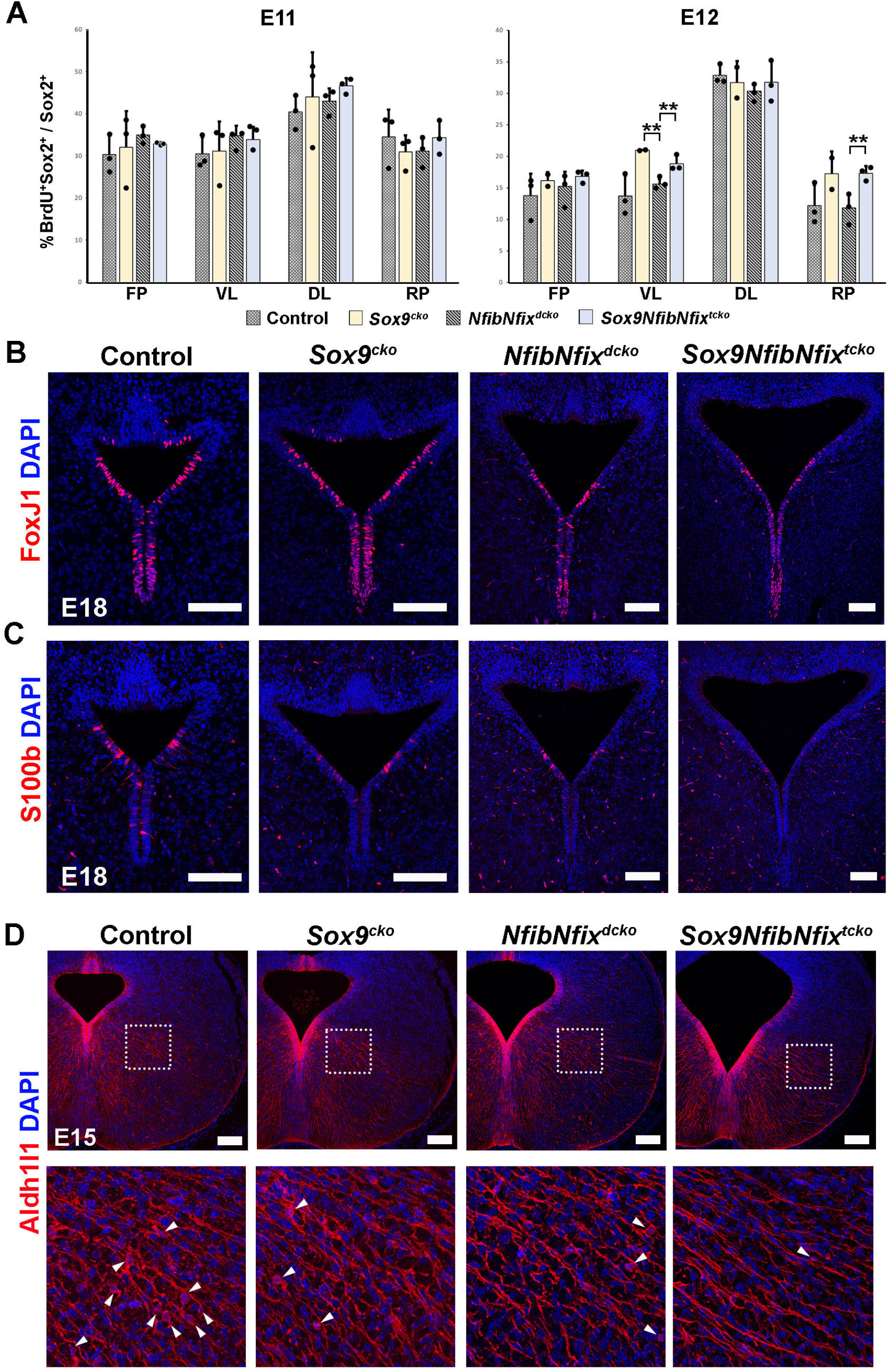
Astrogliogenesis and ependymal maturation in the absence of *Sox9* and *Nfi* factors. **A)** Quantification of 30-minute BrdU uptake in the midbrain at E11 and E12 in different genotypes (n=3). One-way ANOVA with Tukey post hoc test. ** p<0.01. Only comparisons with p-value <0.05 are indicated. **(B-C)** IHC for ependymal markers FoxJ1 and S100b in the E18 VZ of different genotypes, stained on parallel sections. **D)** Aldh1l1^+^ cells in the E15 midbrain of different genotypes. Boxed area indicates the location of the close-ups, with arrowheads pointing to Aldh1l1 signal in the somata of glial cells. Scalebars 100 μm.

**Supplementary Figure 5.**
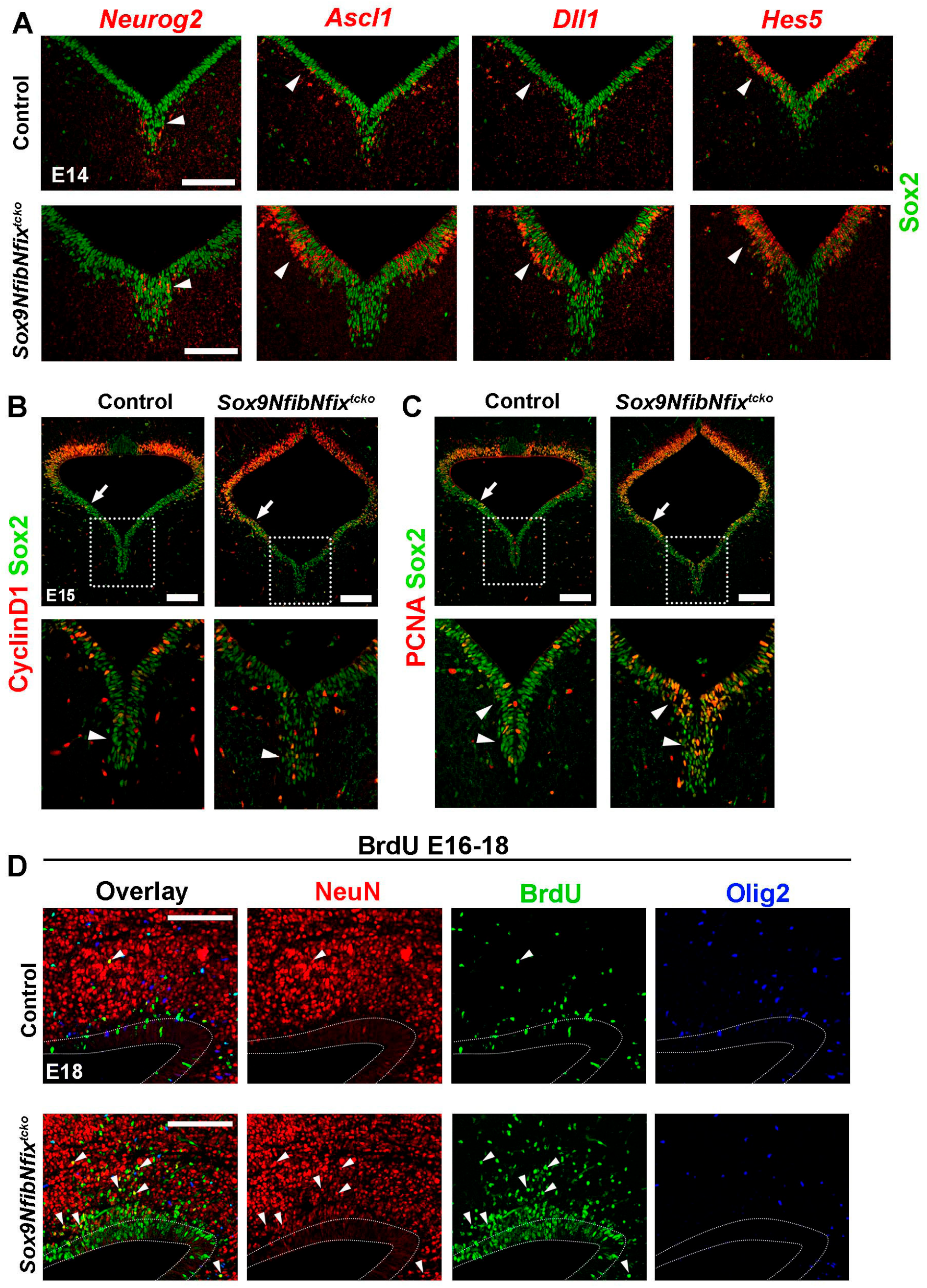
Neurogenic and progenitor marker expression in control and *Sox9NfibNfix*^*tcko*^ midbrain. **A)** Fluorescent in situ hybridization of markers indicated in the E14 midbrain VZ. **(B-C)** IHC detection of CyclinD1 and PCNA in the midbrain VZ, with arrowheads indicating areas where upregulation of signal could be detected in the mutants. Boxed area indicates the location of the close-up image. Regions where increased signal could be seen in the mutants are indicated with arrows and arrowheads. **D)** IHC staining for NeuN, BrdU and Olig2 in the dorsal midbrain in embryos that received continuous BrdU E16-E18. Boundaries of VZ are visualised with dotted lines. Arrowheads point to BrdU+ NeuN+ cells outside the VZ. Scalebars are 200 μm.

**Supplementary Figure 6.**
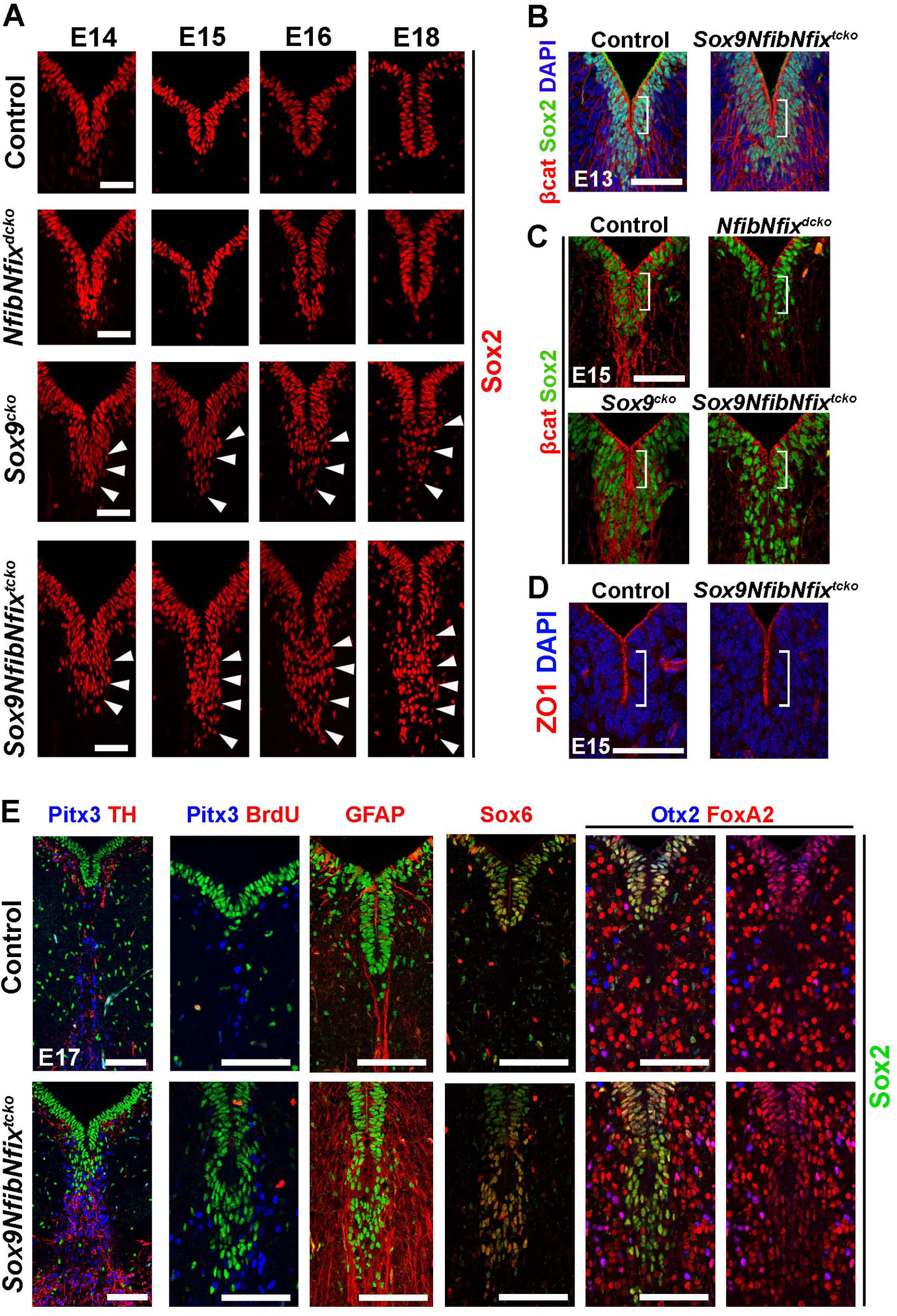
Characterization of ectopic DA progenitors in mutant embryos. **A)** Close-up views if the ventral midbrain VZ in E14-E18 controls and mutants, with arrowheads indicating the ectopic Sox2^+^ cells below the ventricular layer. **B-D)** IHC detection of an adherens junction marker betacatenin and a tight junction marker ZO1 in the ventral midbrain. Brackets indicate the apical layer of the ventralmost VZ. **E)** IHC characterization of the ectopic progenitors in *Sox9NfibNfix*^*tcko*^ at E17 which had received a 3-hour BrdU pulse before tissue collection. Scalebars 50 μm in A-D and 100 μm in E.

**Supplementary Figure 7.**
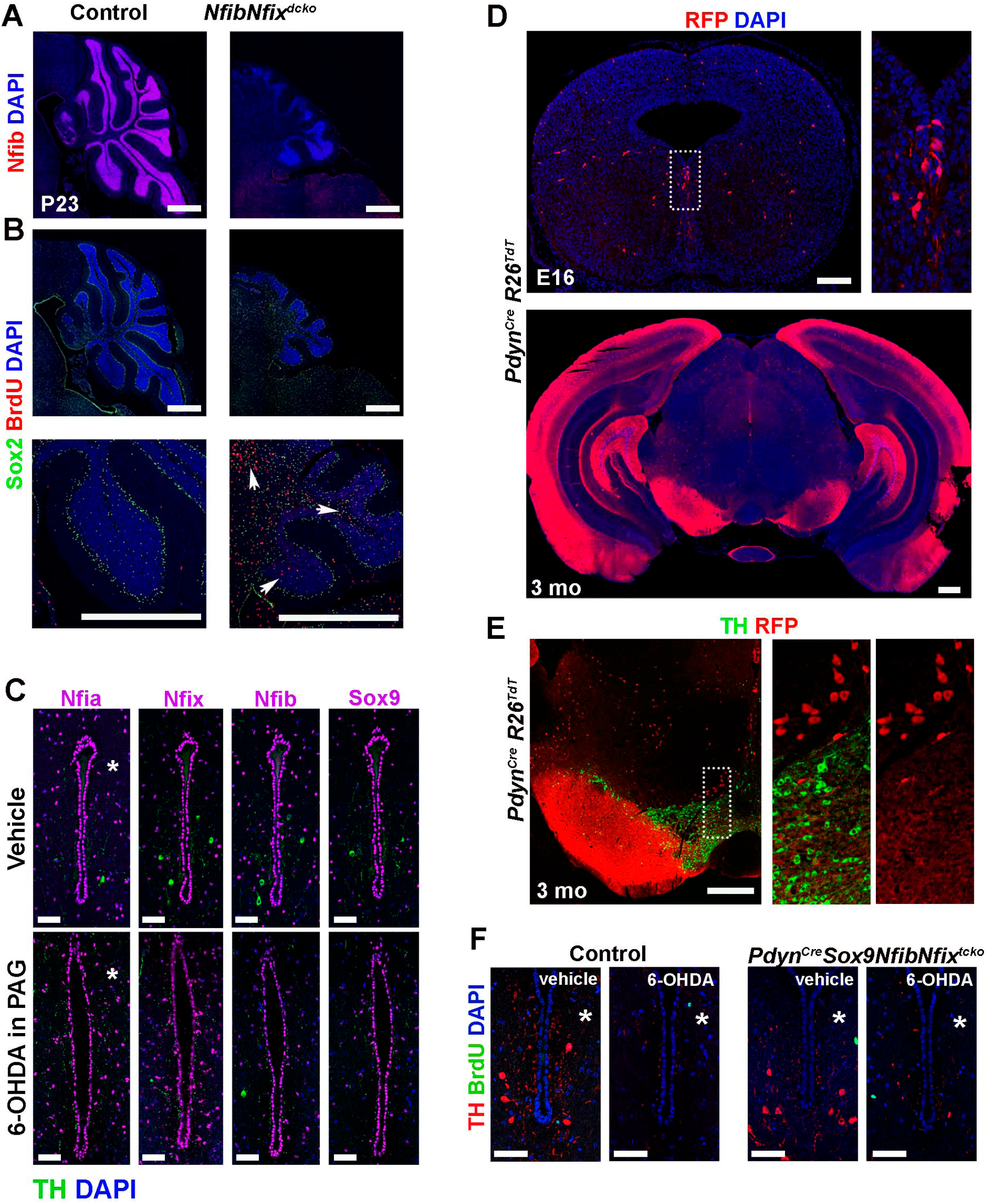
Characterization of adult *Pdyn*^*Cre*^ *Sox9NfibNfix*^*tcko*^ mutants. **A)** Immunohistochemical staining for Nfib in the sagittal sections of cerebellum in control and *Nfib*^*cko*^ animals. **B)** Numerous BrdU^+^ cells were detected in the mutant cerebellum after a 3-day treatment with BrdU in drinking water before tissue collection (arrows). **C)** The expression of Nfi factors and Sox9 in the aqueduct after a local 6-OHDA lesioning of DA neurons in periaqueductal grey. **D)** R26^TdT^ staining pattern in *Pdyn*^*Cre*^ embryonic and adult midbrain. First RFP^+^ cells in the midbrain ventricular layer were detected at E16. In the adult, strong RFP^+^ signal could be seen in the neuronal fibers of SNpr, dorsal and ventral aqueduct, and in parts of hippocampus and cortex. **E)** TH and RFP costaining on adult *Pdyn*^*Cre*^ reporter showing that dopaminergic cells were not labelled with Pdyn^Cre^. **F)** BrdU-labelling in control and *Pdyn*^*Cre*^*Sox9NfibNfix*^*tck*o^ animals after a 6-OHDA-mediated lesioning of DA neurons in the PAG, followed by BrdU in drinking water for 25 days. Scalebars 500 μm in A, B, in 3-month-old brain in D, and E, 200 μm in E16 brain in D, and 50 μm in C and F. The asterisks (*) indicate the injection side.

**Supplementary Figure 8.**
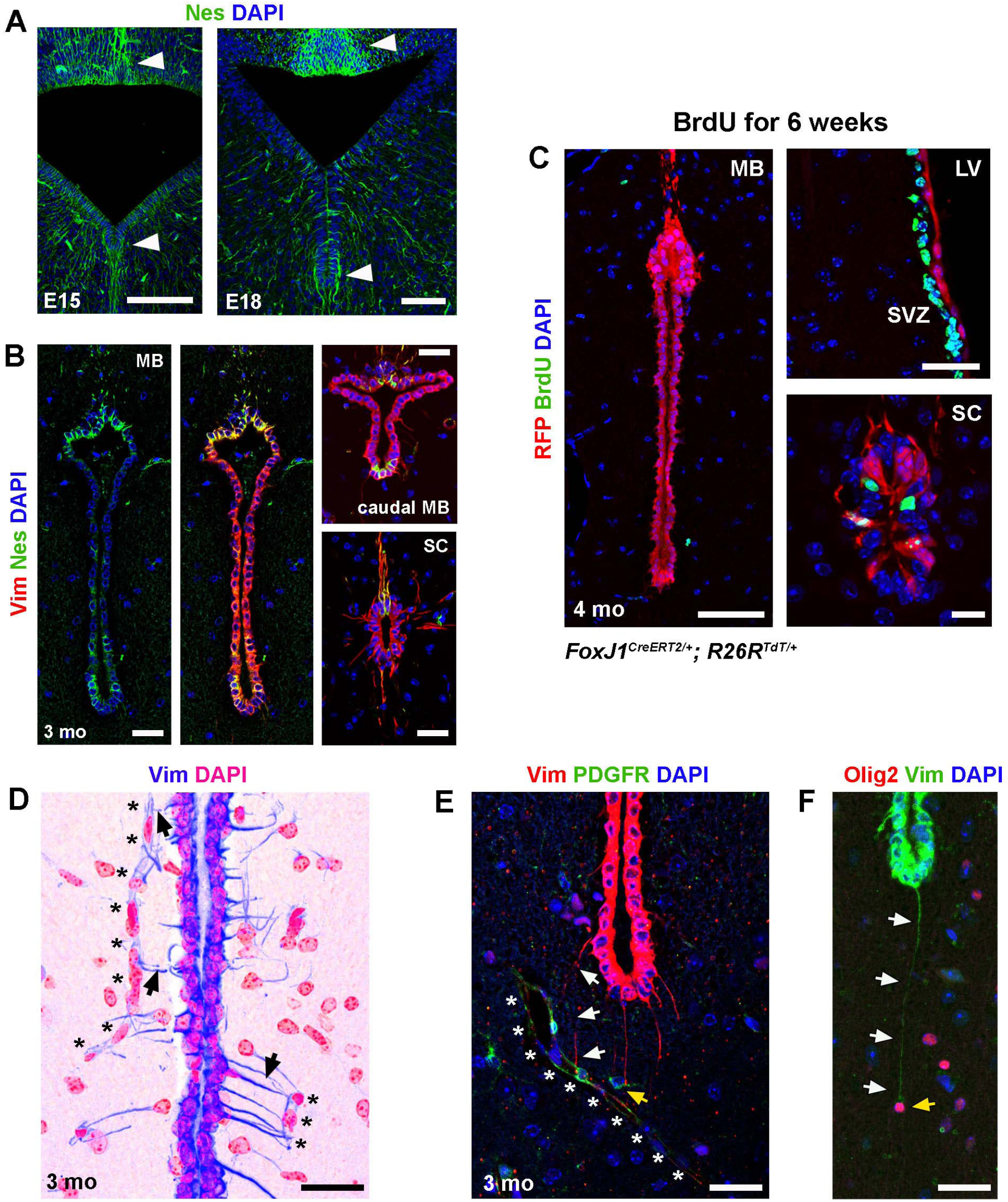
Characterization of midbrain ependymal cells. **A-B)** An IHC staining of Nestin in E15, E18 and in adult showing that the most ventral and dorsal regions of midbrain VZ retain higher expression levels of Nestin compared to the surrounding regions. The most ventral and dorsal tip of adult aqueduct show higher Vimentin and Nestin expression, which resembles the expression pattern in the spinal cord (SC). **C)** Ependymal cells in the adult midbrain do not take up BrdU unlike ependymal cells in the spinal cord or neural stem cells in the subventricular zone of lateral ventricles (LV) after 6 weeks treatment with BrdU in drinking water. **D)** Lateral wall of the adult aqueduct, with inverted pseudocolours for vimentin and DAPI. Several Vim^+^ fibers (arrows) contact nearby blood vessels (indicated by *). **E-F)** Some ventral Vim^+^ fibers (arrows) make contact with blood vessels (asterisks) and PDGFR^+^ or Olig2^+^ cells (yellow arrows). Scalebars 200 μm in A, 100 μm in B, 50 μm in the midbrain image in B, 25 μm in B, D-F, in LV image in C, and 15 μm in the SC image in C.

## Notes

### Competing Interest Statement

The authors have declared no competing interest.

